# RNA and DNA G-quadruplexes bind to human Dicer and inhibit its activity

**DOI:** 10.1101/2020.05.05.078592

**Authors:** Natalia Koralewska, Agnieszka Szczepanska, Kinga Ciechanowska, Marta Wojnicka, Maria Pokornowska, Marek C. Milewski, Dorota Gudanis, Daniel Baranowski, Chandran Nithin, Janusz M. Bujnicki, Zofia Gdaniec, Marek Figlerowicz, Anna Kurzynska-Kokorniak

**Author notes:** Corresponding author: Anna Kurzynska-Kokorniak, Noskowskiego 12/14, 61-704 Poznan, Poland. Contributed equally (Natalia Koralewska, Agnieszka Szczepanska).

## Abstract

Guanine (G)-rich single-stranded nucleic acids can adopt G-quadruplex structures. Accumulating evidence indicates that G-quadruplexes serve important regulatory roles in fundamental biological processes such as DNA replication, transcription, and translation, while aberrant G-quadruplex formation is linked to genome instability and cancer. Understanding the biological functions played by G-quadruplexes requires detailed knowledge of their protein interactome. Here, we report that both RNA and DNA G-quadruplexes are bound by human Dicer *in vitro*. Using *in vitro* binding assays, mutation studies, and computational modeling we demonstrate that G-quadruplexes can interact with the Platform-PAZ-Connector helix cassette of Dicer, the region responsible for anchoring microRNA precursors (pre-miRNAs). Consequently, we show that G-quadruplexes efficiently and stably inhibit the cleavage of pre-miRNA by Dicer. Our data highlight the potential of human Dicer for binding of G-quadruplexes and allow us to propose a G-quadruplex-driven sequestration mechanism of Dicer regulation.

## Introduction

Dicer belongs to the ribonuclease III (RNase III) family of double-stranded RNA (dsRNA)-specific endoribonucleases that are essential for the maturation and decay of coding and noncoding RNAs in both prokaryotes and eukaryotes (1). Proteins classified into this family share a unique fold (RNase III domain) that dimerizes to form a catalytic center where the hydrolysis of phosphodiester bonds occurs, leaving a characteristic dsRNA product with a 5′ phosphate and a 2-nucleotide (nt) overhang with a hydroxyl group at the 3′ end (reviewed in (1)). Human Dicer (hDicer), consisting of 1,992 amino acids (220-kDa), is one of the most structurally complex members of the RNase III family. It comprises an amino (N)-terminal putative helicase domain, a domain of unknown function (DUF283), Platform, Piwi-Argonaute-Zwille (PAZ) domain, a Connector helix, two RNase III domains (RNase IIIa and RNase IIIb) and a dsRNA-binding domain (dsRBD) (2, 3).

Extensive studies have deciphered the roles of the Dicer domains in binding and processing of its canonical substrates, i.e., dsRNAs and single-stranded hairpin precursors of microRNAs (pre-miRNAs). The putative helicase domain selectively interacts with the apical loop of pre-miRNA and helps to discriminate between substrates (4). The DUF283 domain has been implicated in the binding of single-stranded nucleic acids (5), and may therefore be involved in interactions with the apical loop of pre-miRNA hairpins (6). Two adjacent domains, Platform and PAZ, anchor the 5′ phosphate and 2-nt 3′ overhang of a substrate (3). The RNase IIIa and RNase IIIb domains form a single dsRNA-cleavage center (7). Finally, the C-terminal dsRBD is presumed to play an auxiliary role in RNA binding (8).

G-quadruplexes are noncanonical structures formed by guanine-rich DNA and RNA molecules. They are organized in stacks of two or more G-quartets, in which four guanines are associated through Hoogsteen base pairing and stabilized by a monovalent cation, such as Na^+^ or K^+^ (for reviews see (9)). G-quadruplex-forming motifs occur in the genomes and transcriptomes of many species, including humans. Biologically relevant G-quadruplexes were first discovered in eukaryotic telomeres (10). Since then, multiple studies have shown the importance of G-quadruplexes in the regulation of DNA replication, gene expression, and telomere maintenance, and have linked G-quadruplexes to several human diseases (for review see (11)). The formation of G-quadruplexes has also been reported in the regulation of RNA metabolism, including the miRNA pathway, where such structures may influence Dicer activity and other steps of miRNA biogenesis (12-14).

Previously, we characterized the inhibitory potential of structurally diverse short RNA molecules, 12-to 60-nt in length, able to influence processing of pre-miRNA by hDicer either by binding to the enzyme or by base-pairing with the substrate (pre-miRNA) (15-17). We showed that 12-mers are too short to bind to hDicer efficiently, and consequently, to act as competitive inhibitors. However, if these 12-mers base-pair with the apical region of a pre-miRNA hairpin, they preclude the cleavage by hDicer, thereby acting as pre-miRNA-specific inhibitors (16). Interestingly, in contrast to the other tested 12-mers, one guanine-rich molecule bound to hDicer and inhibited cleavage of all tested pre-miRNAs (16). However, our previous studies did not explain the mechanism of this phenomenon.

Here, we demonstrate that a hDicer-binding guanine-rich 12-mer can form a G-quadruplex structure. This finding inspired us to investigate possible interactions between hDicer and nucleic acids adopting G-quadruplex structures. In our studies, we used a full-length hDicer, as well as the hDicer fragment responsible for canonical substrate binding, encompassing the Platform and PAZ domains. We also analyzed several structurally well-characterized short telomeric RNA and DNA G-quadruplexes from human and the ciliate *Oxytricha nova*. Using molecular modeling and mutation studies, we demonstrated that the Platform and PAZ domains can bind both RNA and DNA G-quadruplexes. We showed that binding of RNA and DNA G-quadruplexes by hDicer precludes pre-miRNA cleavage, which suggests the existence of yet another mechanism regulating Dicer activity and miRNA biogenesis *in vivo*.

## Results

### hDicer binds RNA G-quadruplexes

The 12-nt guanine-rich RNA, found to bind to hDicer, was perfectly complementary to the apical loop of pre-mir-210, and consequently, it was named “AL-210” (16). We hypothesized that the structure adopted by AL-210 determined its ability to bind to hDicer and to inhibit its cleavage activity. Accordingly, we used Fold and bifold algorithms provided by the RNAstructure web server (18) to predict the lowest free energy secondary structures for AL-210 and three other 12-mers used in our previous studies: AL-16-1, AL-21, AL-33a (i.e., the oligomers designed to target the apical loops of pre-mir-16-1, pre-mir-21, pre-mir-33a, respectively). The results indicated that AL-33a and AL-210 can form stable homodimers (Supplementary Fig. S1). AL-16-1 and AL-21 were predicted to be monomeric. Since the structure adopted by RNA may depend on the RNA concentration, each 12-nt RNA was assayed by polyacrylamide gel electrophoresis (PAGE), under nondenaturing conditions, at four different concentrations (0.01, 0.1, 1 and 10 µM). Under the applied conditions AL-16-1 and AL-21 always migrated in the gel as a single conformer, whereas AL-33a and AL-210, depending on the concentration, migrated as faster- or slower-moving conformers. The faster-moving conformers seemed to represent monomers (single-stranded RNA, ssRNA) (Fig. 1A). At concentrations equal to or greater than 0.1 µM, AL-33a migrated as the slower-moving conformer, presumably corresponding to a homodimer (dsRNA) form. Interestingly, at a concentration equal to or greater than 1 µM, AL-210 migrated even slower than a putative dsRNA form (Fig. 1A and Supplementary Fig. S1).

**Fig. 1.**
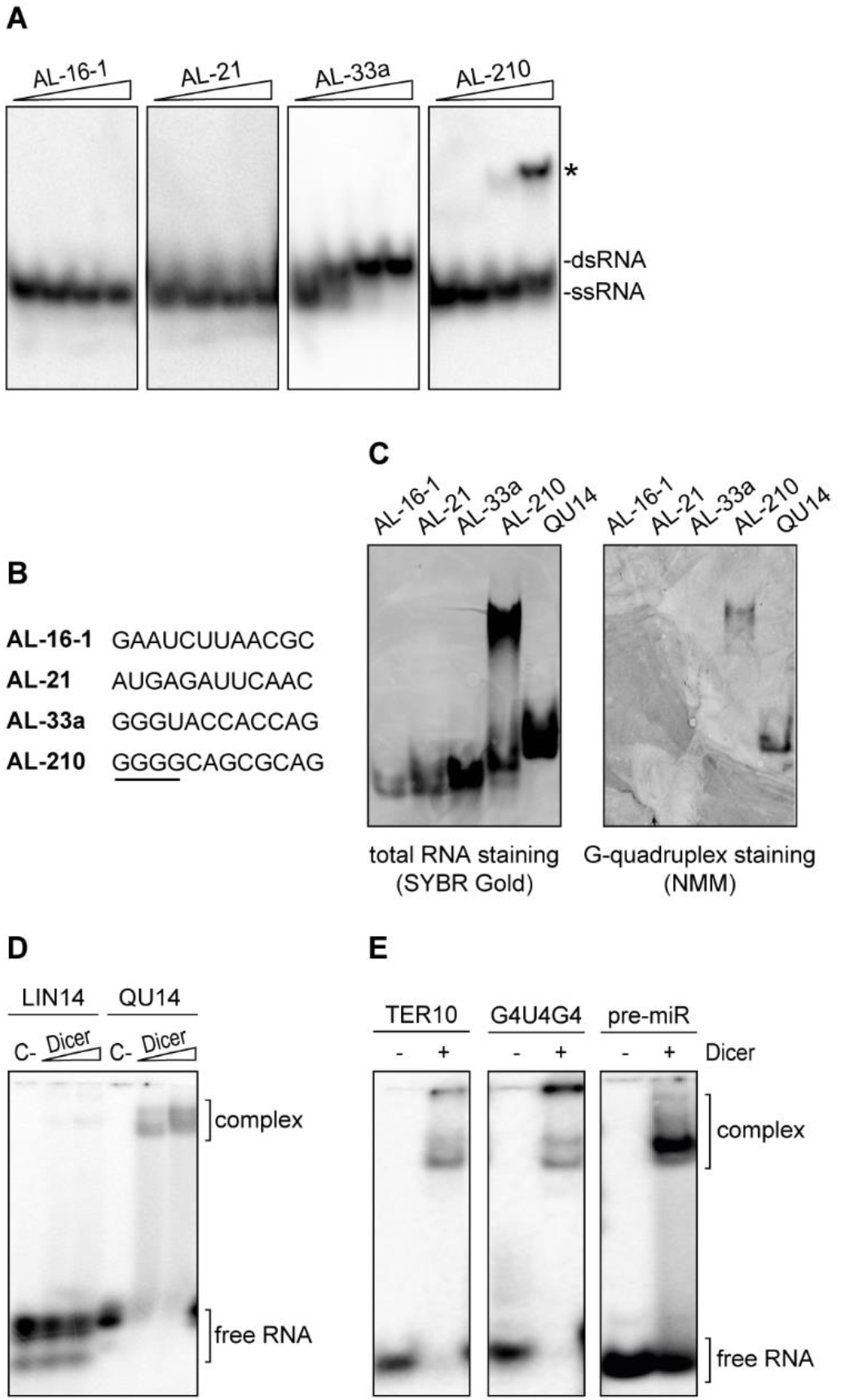
hDicer binds RNA G-quadruplexes. **(A)** Native PAGE analysis of the mixtures of unlabeled and 5′-^32^P-labeled RNA 12-mers (AL). Triangles represent increasing amounts of a given oligomer (0.1, 1, 10 µM). Bands corresponding to single- and double-stranded forms of the oligomers are indicated. **(B)** Comparison of the sequences of RNA 12-mers used in the study. A guanine-tract in the sequence of AL-210 is underlined. **(C)** Native PAGE analysis of RNA 12-mers (AL) and a control 14-nt G-quadruplex (QU14) (100 pmol each). Gels were treated with nucleic acid stain SYBR Gold (*left*) or G-quadruplex-specific dye, NMM (*right*). **(D)** EMSAs with hDicer and the 5′-^32^P-labeled 14-nt RNA not adopting a G-quadruplex structure (LIN14), or the 5′-^32^P-labeled RNA G-quadruplex (QU14). *C-* a control sample with no protein. Triangles represent increasing amounts of hDicer (100, 250 nM). **(E)** EMSAs with hDicer (250 nM) and the 5′-^32^P-labeled RNA G-quadruplex (TER10 or G4U4G4) or a control pre-miRNA. +/- refers to Dicer presence in the reaction mixture.

When analyzing the primary structure of the 12-mers (Fig. 1B), we noticed that a striking feature of AL-210 is a stretch of four guanines at the 5′ end. Such a sequence motif is commonly found in nucleic acids adopting G-quadruplex structures. To test the possibility that the slower-migrating conformer of AL-210 represents a G-quadruplex form, we performed native PAGE followed by G-quadruplex-specific staining with the *N*-methyl mesoporphyrin IX (NMM) fluorescent probe (19). To promote and stabilize a potential G-quadruplex structure, oligomers were incubated in buffer containing 100 mM KCl before the electrophoresis. As a positive control, we used a 14-nt RNA (called “QU14”) known to adopt a G-quadruplex architecture (20). Under the applied conditions, QU14, AL-16-1, AL-21, and AL-33a each migrated as a single conformer (Fig. 1C, left panel), whereas AL-210 migrated as faster- and slower-moving conformers, as demonstrated above (Fig. 1A). Staining with NMM showed that the G-quadruplex-specific dye was bound only by QU14 and the slower-moving AL-210 conformer, but not by AL-16-1, AL-21, AL-33a or fast-moving AL-210 conformers (Fig. 1C, right panel). Together, these results suggest that AL-210 adopts a G-quadruplex structure and, presumably, in such a form, it could bind to hDicer.

We hypothesized that hDicer could also bind other RNAs containing G-quadruplex structures, including QU14. We performed electrophoretic mobility shift assays (EMSAs) involving hDicer and 5′-^32^P-labeled QU14 or a control 14-nt RNA (called “LIN14”) that did not adopt a G-quadruplex structure. QU14 was bound efficiently by hDicer, while only residual binding of LIN14 was detected (Fig. 1D). Weak binding of LIN14 agrees with our previous observations that hDicer does not interact efficiently with RNAs shorter than 20-nt (17). Next, we tested whether hDicer would also bind other short RNAs (< 20-nt) with G-quadruplex structures. Accordingly, we performed EMSAs with hDicer and 5′-^32^P-labeled oligonucleotides with sequences corresponding to the well-characterized telomeric repeat-containing RNAs (TERRAs) of human (21) or ciliate *Oxytricha nova* (22) origin, i.e., r(GGGUUAGGGU) called “TER10” and r(GGGGUUUUGGGG) called “G4U4G4”, respectively (detailed information about guanine-rich oligomers used in the studies is presented in Table S1). Both TER10 and G4U4G4 were bound by hDicer. A control binding reaction contained the canonical hDicer substrate, 58-nt pre-miRNA (Fig. 1E). These results confirm that hDicer is able to bind RNAs adopting a G-quadruplex structure.

### The PPC cassette of hDicer binds RNA and DNA G-quadruplexes

According to the current knowledge about the mechanism of Dicer action, the initial recognition and anchoring of the substrate occur within the region spanning the Platform-PAZ-Connector helix cassette, called “PPC” (3). The PPC contains two adjacent pockets, a 2-nt 3′-overhang-binding pocket (3′-pocket) within the PAZ domain and a phosphate-binding pocket (5′-pocket) within the Platform and PAZ domains (Fig. 2A) (3, 23). To test whether PPC can also bind RNA G-quadruplexes, we produced PPC in a bacterial expression system (Supplementary Fig. S2A) and then carried out EMSAs with PPC and either 5′-^32^P-labeled r[AGGG(UUAGGG)3] called “TER22”, which is an extended version of TER10, or G4U4G4 (Table S1). PPC formed stable complexes with both RNA oligomers, with a Kd value of ∼7 nM for the PPC and TER22 complex (Fig. 2B), and ∼10 nM for the PPC and G4U4G4 complex (Fig. 2C). For comparison, we also determined binding affinities for PPC and 5′-^32^P-labeled 21-nt RNA (called “LIN21”), 58-nt pre-mir-21, and 19-bp RNA duplexes having 2-nt 3’ overhanging ends (called “dsRNA_OV”) (Supplementary Fig. S2B). The Kd values calculated for PPC and RNA substrates were ∼46 nM for LIN21, ∼205 nM for pre-mir-21 and ∼50 nM for dsRNA_OV (Supplementary Fig. S2C). The results indicate that the PPC cassette of hDicer binds RNA G-quadruplexes with higher affinity than it binds a pre-miRNA substrate, a miRNA-size RNA or a miRNA-like duplex.

**Fig. 2.**
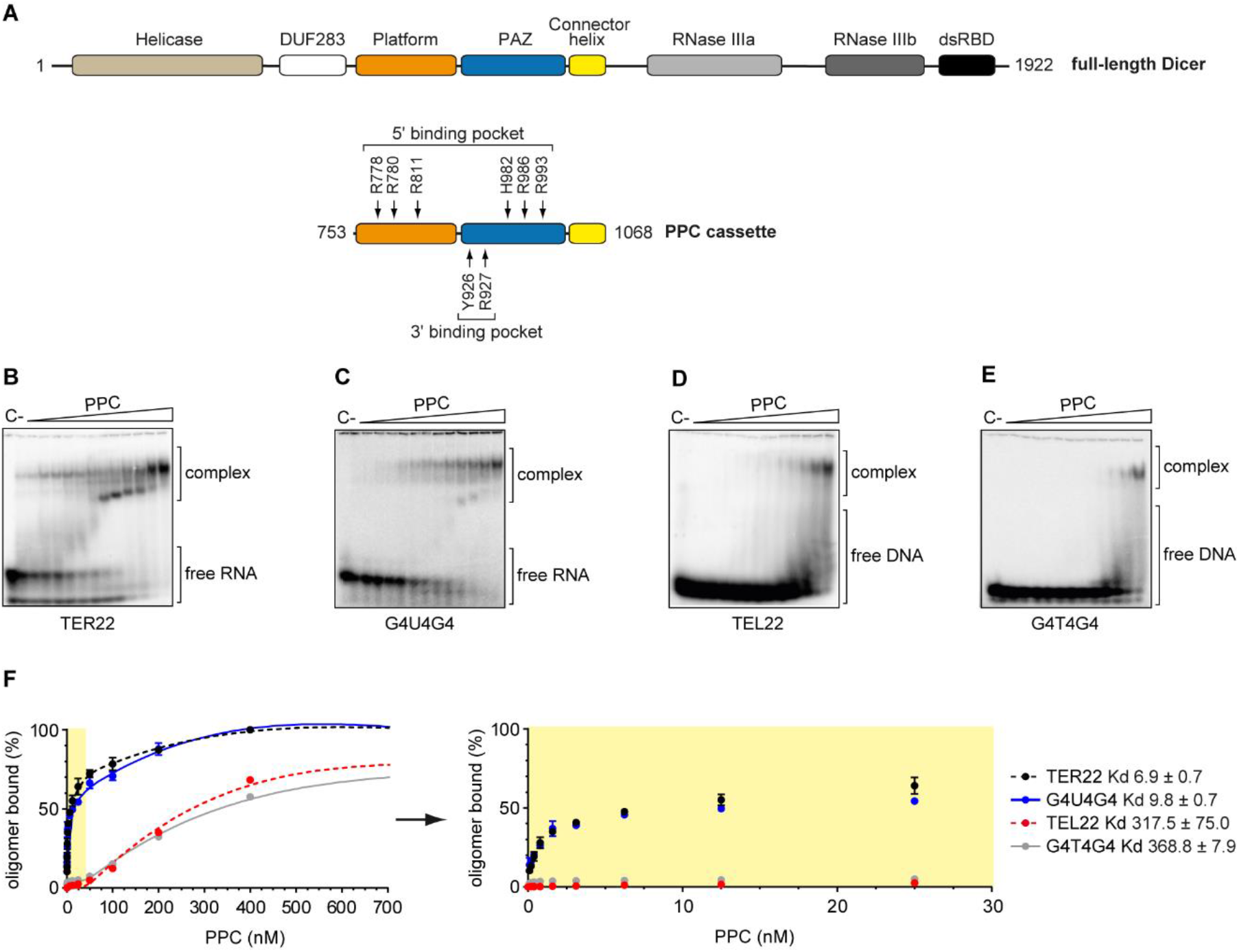
The PPC cassette of hDicer binds RNA and DNA G-quadruplexes. **(A)** Domain architecture of hDicer (*top*) and PPC cassette (*below*). Arrows indicate the RNA-interacting residues within the 5′- and 3′-pocket. **(B, C)** EMSAs with PPC and the 5′-^32^P-labeled RNA G-quadruplexes TER22 (B) and G4U4G4 (C). *C-* a control sample with no protein. Triangles represent increasing amounts of PPC (0.1, 0.2, 0.4, 0.8, 1.6, 3.125, 6.25, 12.5, 25, 50, 100, 200, 400 nM). The Kd of RNA-PPC complexes was determined from the binding isotherms by curve-fitting using nonlinear regression. Error bars represent SD from three separate experiments. **(D, E)** EMSAs with PPC and the 5′-^32^P-labeled DNA G-quadruplexes TEL22 (D) and G4T4G4 (E). *C-* a control sample with no protein. Triangles represent increasing amounts of PPC (0.1, 0.2, 0.4, 0.8, 1.6, 3.125, 6.25, 12.5, 25, 50, 100, 200, 400 nM). The Kd values of DNA-PPC complexes were determined from the binding isotherms by curve-fitting using nonlinear regression. Error bars represent SD from three separate experiments. **(F)** Quantitative analysis of the binding assay between PPC and 5′-^32^P-labeled RNA/DNA G-quadruplexes. Error bars represent SD from three separate experiments.

Given the fact that telomeric repeat-containing sequences are present not only in telomeric RNA but also within telomeric DNA at the chromosome ends, we asked whether PPC could interact with DNA G-quadruplexes as well. Accordingly, we used 5′-^32^P-labeled DNA oligomers with sequences identical to TER22 and G4U4G4, i.e., “TEL22” and “G4T4G4”, respectively (Fig. 2D, E). The Kd value was ∼318 nM for the PPC and TEL22 complex and ∼369 nM for the PPC and G4T4G4 complex. These data indicated that PPC bound DNA G-quadruplexes much weaker than their RNA G-quadruplex counterparts. Additionally, TER22 and TEL22 were bound by PPC slightly better (∼1.4 and ∼1.2 times) than G4U4G4 and G4T4G4, respectively. Together, these results indicate that both the type and length of nucleic acid adopting a G-quadruplex structure determine its binding affinity to the PPC cassette of hDicer.

### Both the 3′-pocket and the 5′-pocket of hDicer PPC cassette are involved in the binding of G-quadruplexes

To test whether oligonucleotides adopting G-quadruplex structures could, like canonical Dicer substrates, be bound in the 3′-pocket and/or 5′-pocket of the PPC cassette, we performed molecular docking and modeling. For this we used the structure of hDicer PPC cassette (PDB entry 4NGF) (3) and the G-quadruplex structures of the 10-nt human TERRA (TER10) (PDB entry 2M18) (21) or *O. nova* G4T4G4 (PDB entry 1JPQ) (22). TER10 formed a dimer of bimolecular G-quadruplexes (21), whereas G4T4G4 appeared as a bimolecular G-quadruplex (22).

We inspected docked poses of both TER10 and G4T4G4 with hDicer protein for clashes with pre-miRNA binding (for detailed information see Materials and Methods). We selected eight best models (docked poses) for the TER10-PPC complex and six best models for the G4T4G4-PPC complex. In seven out of the eight models selected for the TER10-PPC complex, TER10 was positioned within the 3′-pocket of the PPC cassette. In all these models, the 3′ end of one out of the four RNA strands was anchored in the 3′-pocket (Fig. 3A). In one of the eight models, the 3′ end of TER10 was located in the vicinity of amino acid residues lining up the 5′-pocket (Fig. 3B). Three docked poses obtained for the G4T4G4-PCC complex showed that the G4T4G4 quadruplex bound within the 3′-pocket (Fig. 3C), and the other three poses revealed G4T4G4 located within the 5′-pocket of the PPC cassette (Fig. 3D).

**Fig. 3.**
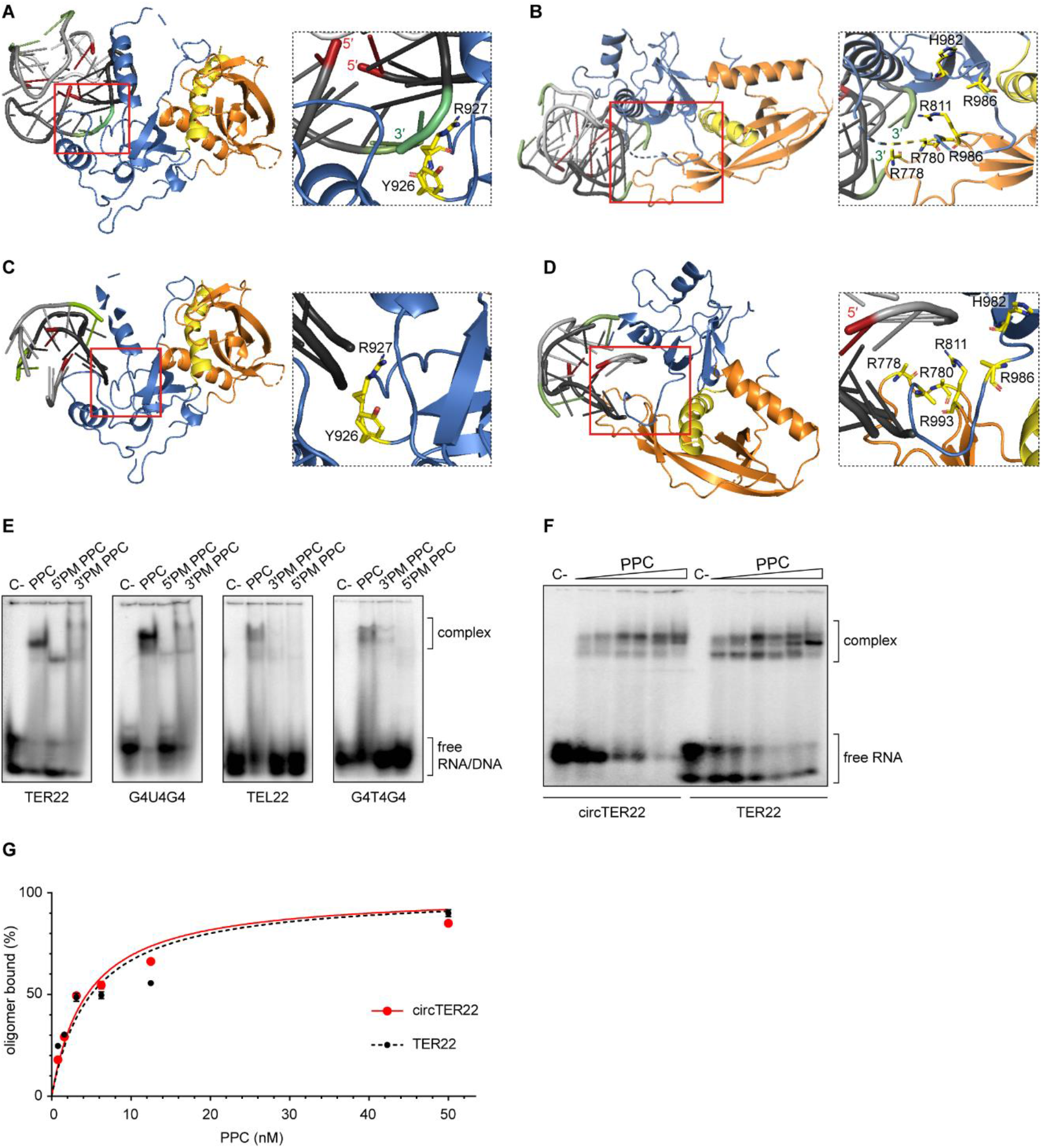
Both the 3′- and 5′-pocket of hDicer PPC cassette are involved in the binding of G-quadruplexes. **(A)** 3D model of TER10 forming a dimer of bimolecular G-quadruplexes bound to the 3′-pocket of hDicer PPC cassette. The 3′- and 5′-ends of the oligonucleotide are colored in green and red, respectively. The Platform is colored in orange, PAZ in blue and Connector helix in yellow. The same color coding is maintained in subpanels B to D of this Fig. **(B)** 3D model of TER10 forming a dimer of bimolecular G-quadruplexes bound to the 5′-pocket of hDicer PPC cassette. **(C)** 3D model of G4T4G4 forming a bimolecular G-quadruplex bound to the 3′-pocket of hDicer PPC cassette. **(D)** 3D model of G4T4G4 forming a bimolecular G-quadruplex bound to the 5′-pocket of hDicer PPC cassette. **(E)** EMSAs with PPC, PPC 5′- and 3′-pocket mutants (5′PM, 3′PM), and 5′-^32^P-labeled G-quadruplexes (TER22, G4U4G4, TEL22, G4T4G4). 500 nM protein and 10,000 cpm (approximately 5 nM) of RNA/DNA were used per lane. *C-* a control sample with no protein. **(F)** EMSAs of binding between PPC and TER22 with free ends (TER22) or TER22 with ligated ends (circTER22). Triangles represent the increasing amount of PPC (0.8, 1.6, 3.125, 6.25, 12.5, 50 nM). *C-* a control sample with no protein. **(G)** Quantitative analysis of the binding assay between PPC and TER22/circTER22 G-quadruplexes. Error bars represent SD from three separate experiments.

To validate these models, we generated two variants of the hDicer PPC cassette, one containing two substitutions in the 3′-pocket (Y926F/R927A variant), and the other containing six substitutions in the 5′-pocket (R778A/R780A/R811A/H982A/R986A/R993A variant) (Supplementary Fig. S3A). These changes significantly affect the binding of small interfering RNA (siRNA) by hDicer PPC cassette (3), and the corresponding full-length hDicer variants have been well characterized biochemically (23). Using EMSAs, we examined the ability of PPC variants to bind 5′-^32^P-labeled TER22, G4U4G4, TEL22, and G4T4G4 oligomers. Changes in both the 3′-pocket and the 5′-pocket reduced PPC binding to all G-quadruplexes analyzed (Fig. 3E). Mutations in the PPC pockets had a greater impact on binding to the DNA G-quadruplexes than to their RNA counterparts. However, this can be explained by the fact that, under the applied reaction conditions, the wild-type protein displayed much lower maximum binding capacity for DNA G-quadruplexes than for the corresponding RNA G-quadruplexes (Fig. 2B-E), and mutations in the PPC pockets only proportionally decreased binding. The lower binding capacity of PPC to DNA G-quadruplexes than to RNA G-quadruplexes may be due to the participation of ribose 2′-hydroxyl groups in hydrogen bonding with amino acid residues of the PPC cassette. Taken together, binding assays conducted for the wild-type protein and the 3′-pocket and the 5′-pocket variants (Fig. 3E) corresponded well with the models of PPC and G-quadruplex complexes (Fig. 3A-D), collectively indicating that both pockets of the hDicer PPC cassette are important for the binding of G-quadruplexes.

To investigate whether the free ends of the RNA adopting a G-quadruplex structure are necessary for its binding to the hDicer PPC cassette, we circularized 5′-^32^P-labeled TER22 using T4 RNA ligase, as described previously (24) (Supplementary Fig. S3B, C). In adequate protein concentrations, PPC bound this “circTER22” with similar efficiency as the TER22 RNA with free ends (Fig. 3F, G), indicating that hDicer PPC cassette binds G-quadruplexes no matter whether their 3′- or 5′ ends are available.

### RNA G-quadruplexes bound by the PPC cassette retain their structure

To test whether RNA G-quadruplexes retain their structures upon binding the hDicer PPC cassette, we conducted a binding assay involving TER22 with a derivative of 3,6-bis(1-methyl-2-vinyl-pyridinium) carbazole diiodide (*o*-BMVC) covalently attached to its 5′-end (Supplementary Fig. S3D). *o*-BMVC is a fluorescent light-up probe that selectively binds to G-quadruplex structures (25). As a control, we used 32-nt RNA (called “LIN32”) that does not contain a G-quadruplex motif, labeled with *o*-BMVC in the same way as TER22. Each oligomer was incubated with PPC, and the reaction mixtures were analyzed in a polyacrylamide gel under native conditions. After electrophoresis, we exposed the gel first to 532 nm light to detect bands corresponding to the RNA species adopting G-quadruplex structures (Fig. 4A). Subsequently, we stained the gel with SYBR Gold solution and exposed it to 473 nm light to visualize the total RNA pool (Fig. 4B). The results showed that RNA G-quadruplexes retained their structure upon binding the PPC cassette.

**Fig. 4.**
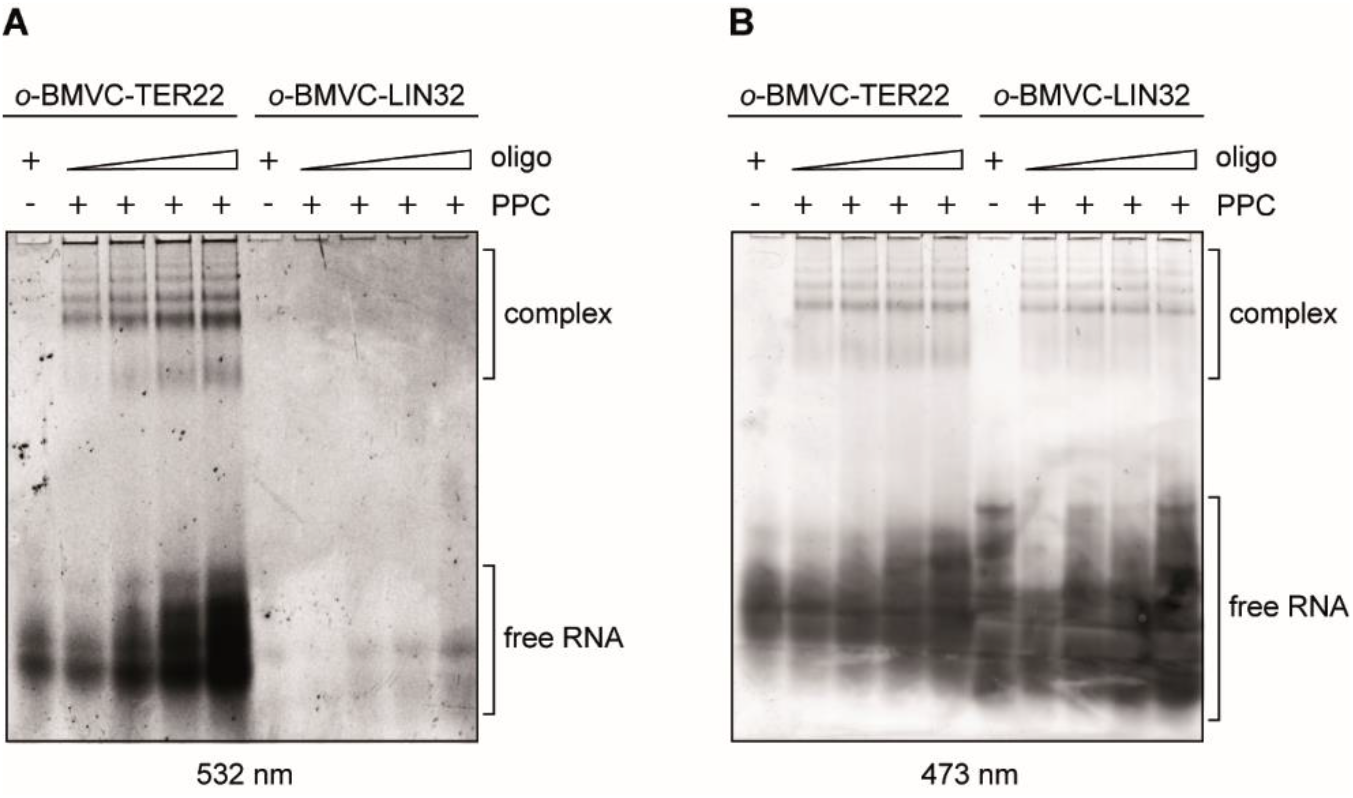
RNA G-quadruplexes bound by the PPC cassette retain their structure. **(A)** EMSAs with 500 nM PPC and TER22 or LIN32, which does not adopt a G-quadruplex structure. Both RNA oligonucleotides were labeled at the 5′-end with a G-quadruplex specific probe (*o*-BMVC). +/- indicates the presence or absence of PPC, or the RNA oligomer. Triangles represent increasing amounts of the RNA oligomer (7.5, 15, 22.5, 30 µM). To visualize RNA adopting G-quadruplex structures, the gel was exposed to 532 nm light. **(B)** The same gel as presented in panel (A) exposed to 473 nm light after staining with SYBR Gold to visualize total RNA.

### RNA and DNA G-quadruplexes inhibit pre-miRNA processing by hDicer

*In vitro*, RNA oligonucleotides bound to hDicer inhibit the cleavage of pre-miRNAs by this enzyme (15, 16). To investigate whether oligonucleotides containing G-quadruplex motifs can also influence the cleavage activity of hDicer, we used RNA oligomers representing human and *O. nova* TERRA (TER10, TER22, G4U4G4) and DNA oligomers representing the corresponding telomeric repeats (TEL22, G4T4G4). As well as those mentioned above, we used three RNAs of 12 to 20-nt: TER12, TER18, TER18-2A. All of the chosen oligomers are known to adopt various G-quadruplex architectures (Table S1). To assess the effect of RNA and DNA G-quadruplexes on hDicer cleavage activity, we performed a set of assays involving hDicer, 5′- ^32^P-labeled pre-mir-21 or pre-mir-33a, and a respective oligomer. The particular pre-miRNAs were chosen as they did not interact with tested oligomers (Supplementary Fig. S4A), which ensures that the effects observed in the experiment does not stem from direct RNA-DNA or RNA-RNA binding, but rather, from protein-DNA or RNA interaction. Moreover, pre-mir-21 and pre-mir-33a represent structurally distinct substrates (Supplementary Fig. S5E).

First, we carried out cleavage assays with RNA G-quadruplexes. The efficiency of pre-miRNA (∼5 nM) cleavage in reactions with individual oligomers applied at one of three concentrations (0.1, 0.5, or 2 µM) was normalized to the cleavage efficiency in a control reaction with no oligomer added. Another set of control reactions involved 12-nt RNA (called “LIN12”), which does not bind to hDicer, nor adopts a G-quadruplex structure. In all cases, upon the addition of an RNA G-quadruplex, we observed a dose-dependent inhibition of the cleavage of pre-mir-21 (Fig. 5A and Supplementary Fig. S4B) and pre-mir-33a (Fig. 5B and Supplementary Fig. S4C). At the lowest concentration of any of the tested RNA G-quadruplexes, the level of miRNA was reduced by at least 50% in comparison to the control reaction with no oligomer added. When the highest concentration of any of the tested RNA G-quadruplexes was applied, the cleavage of pre-miRNA was abolished by at least 80%. No decrease in the miRNA level was observed in control reactions with LIN12. We did not find any apparent correlation between the length or architecture of the RNA G-quadruplexes and the degree of inhibition they exerted.

**Fig. 5.**
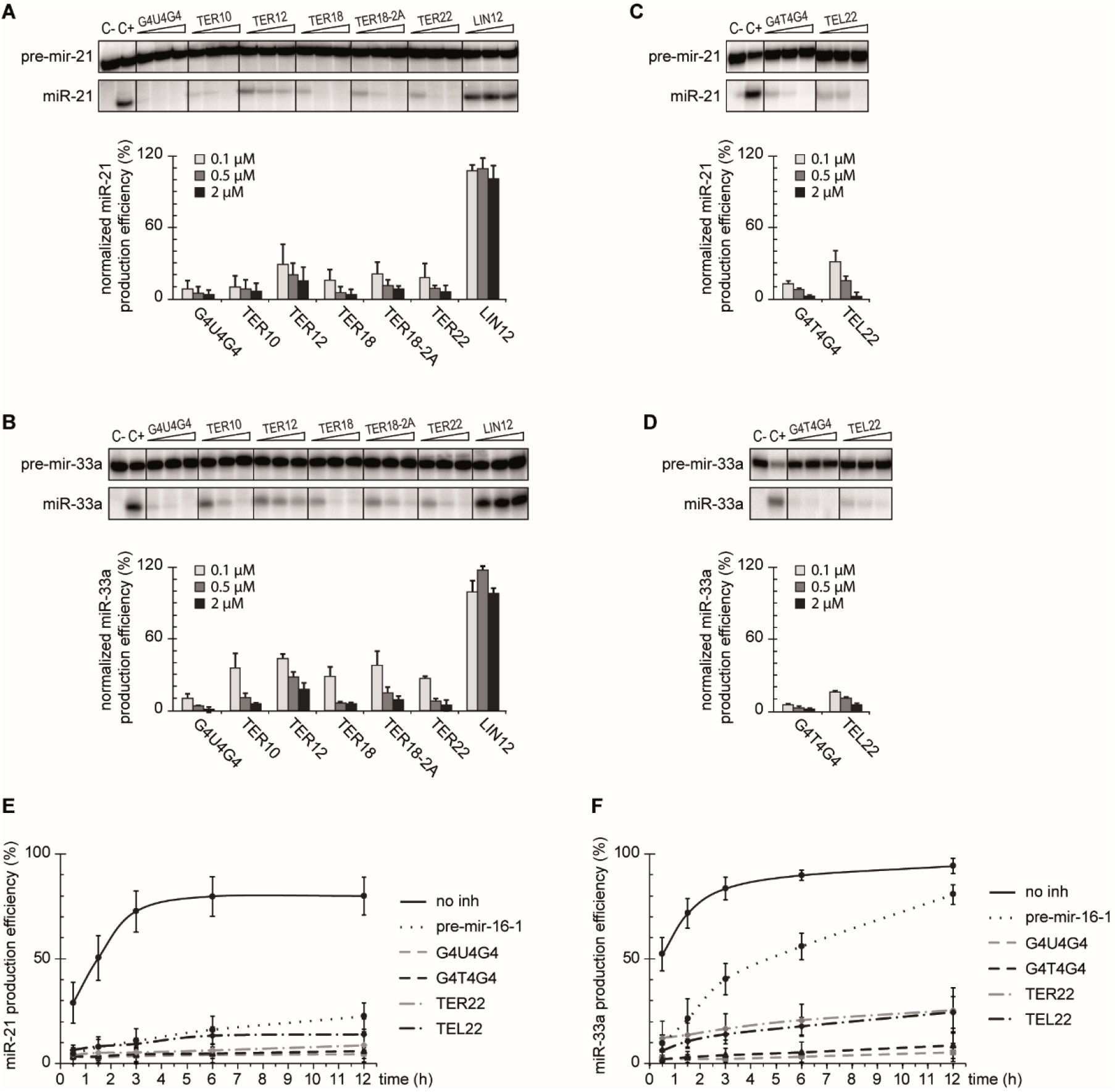
G-quadruplexes inhibit cleavage of pre-miRNA by hDicer in a dose-dependent manner. **(A, B)** Inhibition assay to assess the effect of RNA G-quadruplexes on the cleavage of pre-mir-21 (A) or pre-mir-33a (B) by hDicer. Reactions were carried out for 30 min at 37°C under the low-turnover conditions. LIN12 – a control 12-mer not adopting a G-quadruplex structure, *C-* a sample with no protein, nor inhibitor added, *C+* a sample with hDicer, without inhibitor. Graphs show miRNA production efficiency normalized to the level of miRNA generated in *C+*. Error bars represent SD from three separate experiments. See also Figure S4B,C for full gel images. **(C, D)** The results of the inhibition assay performed similarly as in (A,B) for selected DNA G-quadruplexes and pre-mir-21 (C) or pre-mir-33a (D). See also Figure S4D, E for full gel images. **(E, F)** Quantitative analysis of the time course of hDicer inhibition by G-quadruplexes in reactions with pre-mir-21 (E) or pre-mir-33a (F). Error bars represent SD from three separate experiments. See also Figure S5A-D for representative gel images.

Next, we investigated whether the cleavage of pre-miRNA (∼5 nM) by hDicer can be affected by DNA counterparts of G4U4G4 and TER22, i.e., G4T4G4 and TEL22 (Fig. 5C, D and Supplementary Fig. S4D, E). At the lowest concentration (0.1 µM), G4T4G4 inhibited cleavage of pre-mir-21 and pre-mir-33a by ∼90%, while at the highest concentration (2 µM), the inhibition reached ∼96% for pre-mir-21 and 100% for pre-mir-33a. We observed a similar tendency for TEL22; it inhibited the cleavage of pre-mir-21 and pre-mir-33a by ∼70% at the lowest concentration (0.1 µM), and by ∼95% at the highest concentration (2 µM). The results are comparable with those obtained for G4U4G4 (∼90% and ∼100% inhibition at 0.1 µM and 2 µM concentration, respectively) and TER22 (∼75-80% and ∼95% inhibition at 0.1 µM and 2 µM concentration, respectively), which indicates no differences in the inhibition potency between RNA G-quadruplexes and their DNA G-quadruplex counterparts under the applied reaction conditions (Fig. 5A-D and Supplementary Fig. S4B-E).

Subsequently, we performed a time course assay to measure the level of miR-21 or miR-33a produced in reaction with either no inhibitor, a selected G-quadruplex (G4T4G4, G4U4G4, TEL22, TER22) or pre-mir-16-1 added as a competitor. The reactions were performed under the low-turnover conditions, i.e. a 2-fold molar excess of hDicer over a substrate was used. Additionally, the G-quadruplexes and the competitor were in 50-fold molar excess to hDicer. As expected, in control reactions with either pre-mir-21 or pre-mir-33a and hDicer, but no other oligomer added, we observed a hyperbolic relation between the yield of miRNA and the incubation time (Fig. 5E, F and Supplementary Fig. S5A-D). After 12 h, ∼90% of either of the substrates was processed by the enzyme.

In reactions with pre-mir-21 as a substrate and pre-mir-16-1 as a competitor, we observed a significant inhibition of pre-mir-21 cleavage at all analyzed time points; even after 12 h only ∼22% of pre-mir-21 was processed (Fig. 5E and Supplementary Fig. S5A, B). We also observed stable inhibition for the tested G-quadruplexes; the effect exerted by them was even more prominent than in the case of pre-mir-16-1: after 12 h only ∼5% of pre-mir-21 was processed in reactions with G4T4G4 or G4U4G4, and ∼10% in reactions with TEL22 or TER22 (Fig. 5E and Supplementary Fig. S5A, B).

In reactions with pre-mir-33a as a substrate, and pre-mir-16-1 as a competitor, initially the levels of miR-33a were much lower than in the control reactions without the inhibitor (e.g., ∼10% vs ∼50% after the first 30 min of the reaction) (Fig. 5F and Supplementary Fig. S5C, D) but with time the inhibition was gradually abolished; after 12 h the amount of pre-mir-33a processed in the reaction with the competitor and in the control reaction reached ∼80% and ∼95%, respectively. In reactions with the G-quadruplexes, we also observed low initial levels of miR-33a as in the analogous reactions with pre-mir-16-1 and the reduction of the inhibition with time (Fig. 5F and Supplementary Fig. S5C, D). The amount of pre-mir-33a cut by hDicer increased from ∼5% after 30 min to ∼10% after 12 h in reactions with G4T4G4 or G4U4G4, and from ∼10% to ∼25% in reactions with TEL22 or TER22. Despite the observed accumulation of miRNA over time, and in contrast to pre-mir-16-1, the inhibitory effect exerted by G-quadruplexes remained high even after 12 h of incubation (∼75% decrease in miRNA production in comparison to the control reaction in the case of G-quadruplexes, vs ∼15% in the case of pre-mir-16-1).

Altogether, these findings indicate that RNA and DNA adopting a G-quadruplex structure can affect the cleavage of pre-miRNA by hDicer. Under the low-turnover conditions (excess enzyme to substrate), RNA and DNA G-quadruplexes exerted a similar inhibition effect on the hDicer cleavage of both pre-miRNAs used (Fig. 5E, F). G4U4G4 and G4T4G4 were slightly better inhibitors of pre-mir-21 and pre-mir-33a cleavage than TER22 and TEL22 (95% vs 90% inhibition of pre-mir-21 cleavage after 12 h, and 90% vs 75% inhibition of pre-mir-33a cleavage after 12 h, respectively) (Fig. 5E, F and Supplementary Fig. S5A-D).

Subsequently, we performed a time course assay under the high-turnover conditions using a 50-fold molar excess of a substrate to hDicer, and a 100-fold molar excess of the G-quadruplexes or the competitor. We found that G4U4G4, G4T4G4 and TER22 retained their inhibitory potential after 12 h incubation (Supplementary Fig. S6), whereas the inhibition of either miR-21 or miR-33a production was overcome in reactions with TEL22 (Supplementary Fig. S6). Similar results as for TEL22 were obtained for pre-mir-16-1 competitor, therefore, we conclude that under the applied reaction conditions (1:2 molar ratio of a substrate and an inhibitor), TEL22 can act as a competitive inhibitor. This conclusion is supported by the Kd values calculated for the PPC and TEL22 complex (Kd ∼318 nM), and the PPC and pre-miRNA complex (Kd ∼205 nM). However, considering the binding affinities, we cannot explain why G4T4G4 was not outcompeted by the substrate over the incubation time. TEL22 represents the human telomeric sequence, while G4T4G4 corresponds to ciliate *O. nova* telomeric DNA. Given that the human enzyme was used in the studies, a species-specific regulatory mechanism for Dicer binding to telomeric DNA might be responsible for the effects we observed.

Under the low-turnover conditions, we did observe a difference in the effect of pre-mir-16-1 competitor on pre-mir-21 and pre-mir-33a cleavage by hDicer (Fig. 5E, F). Despite the addition of 100-fold molar excess of pre-mir-16-1 with respect to the other pre-miRNA, after 12 h incubation, ∼80% of pre-mir-33a (Fig. 5F) and only ∼22% of pre-mir-21 were processed (Fig. 5E). These results can be explained by the differences among the structures of the three pre-miRNAs. Pre-mir-21 adopts the most compact structure, with the smallest terminal loop, whereas structures of pre-mir-33a and pre-mir-16-1 contain large internal loops and bulges. In addition, pre-mir-16-1 has the most relaxed terminal loop region (Supplementary Fig. S5E). It has been reported that the pre-miRNA structure influences the efficiency of miRNA processing by Dicer (26). Additionally, the results of our previous studies have indicated that pre-miRNAs can compete for binding to hDicer (15). Based on these data and the models generated for PPC and individual G-quadruplexes (Fig. 3A-D), we propose that a pre-miRNA and a G-quadruplex compete for binding to substrate-anchoring domains of hDicer, i.e., PAZ and Platform. Since the interactions between pre-miRNA and Dicer encompass not only the PPC region of Dicer, but other Dicer domains as well (4, 6) (Fig. 6A), we hypothesize that the competition between two pre-miRNAs for binding to Dicer is more complex, compared with the case involving a pre-miRNA and a G-quadruplex. Consequently, we deduce that under the low-turnover conditions, even despite the high excess of the inhibitor, pre-mir-33a can outcompete pre-mir-16-1 from binding to hDicer, which was not observed for pre-mir-21. However, under the high-turnover conditions, when the excess of a substrate to the enzyme was applied, and the substrate to inhibitor molar ratio was low (1:2), the degree of the miRNA production inhibition caused by pre-mir-16-1 was similar in the case of both substrates (Supplementary Fig. S6E, F).

**Fig. 6.**
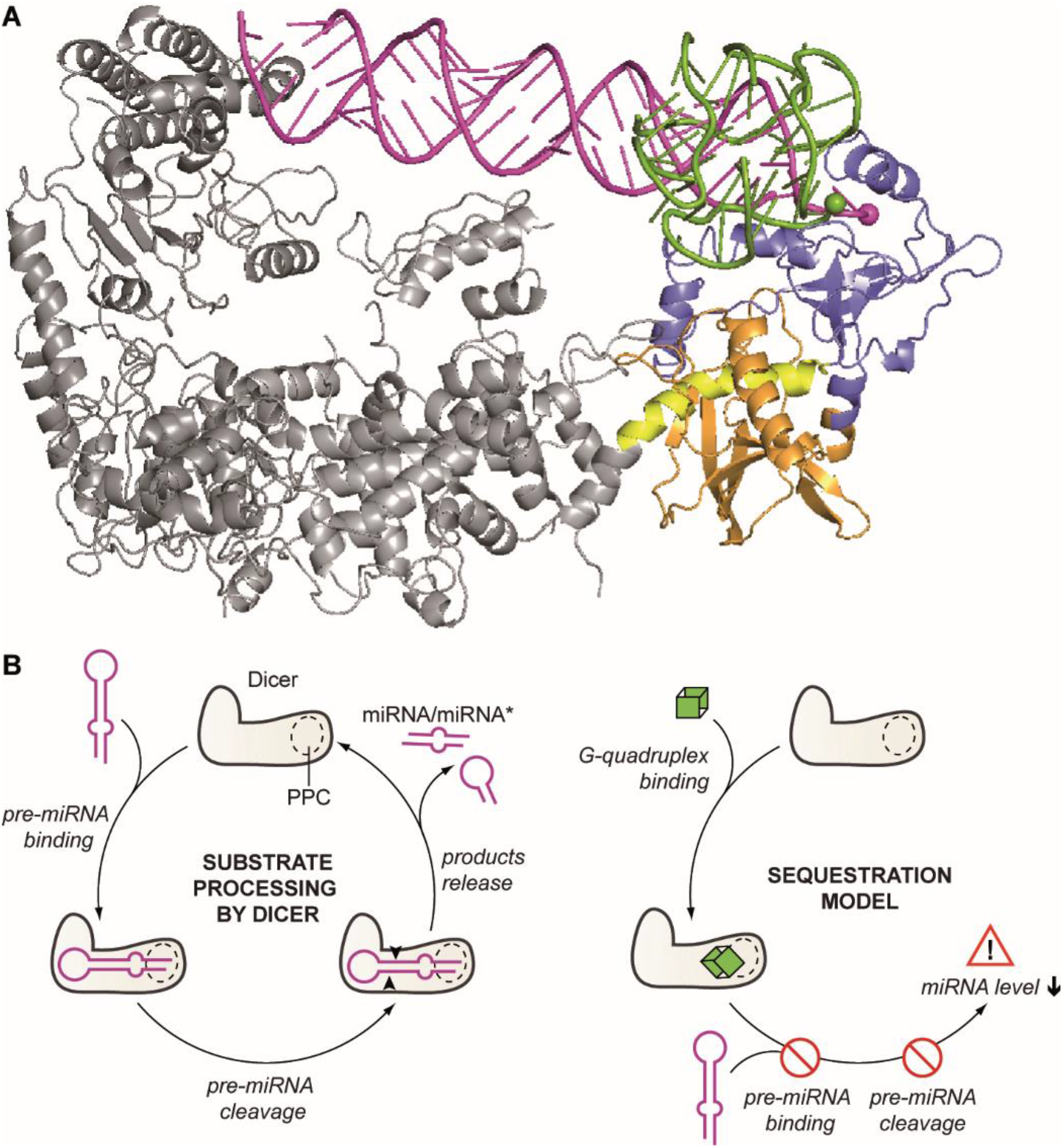
Both pre-miRNA and G-quadruplexes are anchored within the same region of hDicer PPC cassette. **(A)** Superposed structures of hDicer in complex with a pre-miRNA substrate (PDB entry 5ZAL, does not include 15 nt from the apical loop of the pre-miRNA) and PPC cassette in complex with TER10. The Platform is colored in orange, PAZ in blue and Connector helix in yellow, the remaining portion of hDicer is presented in grey; pre-miRNA – in magenta, and TER10 – in green. 3′ ends of pre-miRNA and TER10 within the 3′-binding pocket of PAZ are marked as spheres. **(B)** Putative mechanism of sequestration-dependent regulation of Dicer activity by G-quadruplexes. Dicer anchors pre-miRNA ends within the PPC binding pockets and cleaves precursors to release miRNA products (*left*). G-quadruplexes compete with pre-miRNA for binding to Dicer. The enzyme sequestered in complex with a G-quadruplex cannot bind pre-miRNA and does not generate miRNA (*right*).

## Discussion

Dicer homologs are widely known for their important role in the biogenesis of small regulatory RNAs: miRNAs and siRNAs. However, growing evidence shows that, apart from their canonical role, Dicer proteins serve a number of other functions (for review see (27)). As an endoribonuclease, Dicer participates in processing of diverse groups of RNA, e.g., tRNA (28), DNA-damage-induced RNA (29) or transposable elements (30). Additionally, in *Caenorhabditis elegans*, a caspase-cleaved form of Dicer acts as a deoxyribonuclease that introduces breaks in chromosomal DNA during apoptosis (31). Moreover, the emerging evidence points to possible cleavage-independent regulatory roles of Dicer; in *C. elegans* and human cells, Dicer binds various RNAs, including mRNAs and long non-coding RNAs, passively, i.e., without further cleavage (32). Furthermore, hDicer can act as a nucleic acid annealer, by facilitating base pairing between two complementary RNAs or DNAs (5, 33). Our finding that hDicer can bind RNA and DNA G-quadruplexes, and that this interaction influences Dicer ability to process its canonical substrates, opens new avenues of research on the cellular functions of Dicer.

A variety of proteins bind G-quadruplex-forming RNAs (34). In most cases, however, it is not clear whether the binding involves the G-quadruplex structure or a guanine-rich RNA sequence. In the current study, we demonstrated that G-quadruplexes do not constitute a steric hindrance for hDicer binding (Fig. 1D-E). We also showed that the PPC cassette of hDicer binds RNA G-quadruplexes (Fig. 2B, C) with much higher affinity than it binds non-G-quadruplex RNAs (Supplementary Fig. S2B). Based on the models of the PPC and G-quadruplex complexes (Fig. 3A-D), and mutation studies (Fig. 3E), we propose that RNA and DNA molecules that adopt G-quadruplex structures can bind to either the 3′-pocket or the 5′-pocket of the hDicer PPC cassette. Although the results of our computational modeling indicated that binding of an RNA G-quadruplex by PPC may involve anchoring the 3′ end of the oligonucleotide (Fig. 3A, B), we found that a 5′–3′-end-ligated G-quadruplex can be bound by PPC as well (Fig. 3F). Future work will reveal whether Dicer binds G-quadruplex-containing nucleic acids in cells and what the biological significance of this activity is.

Our results indicate that hDicer can bind both RNA and DNA G-quadruplexes. A search of the transcriptome-wide map of hDicer targets (32), using Quadron software (35), revealed that Dicer binds to several guanine-rich RNAs (Dataset S1). We can speculate about the possible implications of interactions between Dicer and an RNA G-quadruplex. For example, G-quadruplexes may act as molecular anchors that bind and sequester Dicer. A similar role has already been proposed for another type of RNA structure recognized by Dicer, i.e., RNA hairpins formed within endogenous transcripts (32) or adopted by viral RNAs (36). Short RNAs, including miRNAs (37), piRNAs (38), and tRNA fragments (39, 40), can adopt G-quadruplex structures. The length of these RNAs (∼20-30 nt) is close to the length of the guanine-rich oligonucleotides used in the present study (10-22 nt). Accordingly, we hypothesize that short RNAs adopting G-quadruplex structures may sequester Dicer *in vivo*. Such a mechanism could be a part of autoregulatory loops, in which Dicer-processed guanine-rich miRNAs, in the form of G-quadruplexes, act in a negative feedback regulation of Dicer (17, 41). The sequestration of Dicer by RNA G-quadruplexes may control the pool of Dicer available for the biogenesis of miRNA. This hypothesis is supported by the results of the inhibition assay presented in Fig. 5, Supplementary Fig. S5A-D and S6. The observed RNA G-quadruplex-driven inhibition of pre-miRNA cleavage by hDicer was very efficient and stable over time. Since, under the applied reaction conditions, the binding affinity of hDicer PPC for the RNA G-quadruplex (Kd ∼7-10 nM) (Fig. 2B, C) was much higher than the binding affinity of hDicer PPC for the pre-miRNA (Kd ∼205 nM) (Supplementary Fig. S2B), we conclude that an RNA G-quadruplex, once bound to hDicer, occupies the substrate binding pockets of PPC thereby blocking access for a pre-miRNA (Fig. 6B). This is in contrast to a pre-miRNA substrate that, once bound and cleaved by hDicer, leaves the binding pockets, making them available for another substrate molecule.

Proteins binding RNA G-quadruplexes are important for the intracellular transport of mRNA for its local translation (42). Moreover, they act as a switch that controls the access of miRNA to its mRNA targets embedded in guanine-rich regions adopting G-quadruplex structures (43, 44). Since Dicer binds both miRNA (32) and RNA G-quadruplexes, it may be directly involved in the control of translation of mRNAs that form local G-quadruplex structures. Possible direct involvement of Dicer in the posttranscriptional control of gene expression has been already extensively investigated in the context of interactions between miRNA-bound Dicer and complementary mRNA targets (33).

Dicer has been detected in the nucleus (45), which together with our finding that hDicer binds to DNA G-quadruplexes, leads us to propose that Dicer may interact with telomeric DNA G-quadruplexes or be involved in the transcriptional regulation of gene expression by binding to G-quadruplexes in promoter regions.

Altogether, we have demonstrated that G-quadruplexes can bind to Dicer and affect its activity. The results reported here support the notion that Dicer is a versatile protein, whose function is not limited to RNase III activity.

## Materials and Methods

### Oligonucleotides

Sequences of all oligonucleotides used in this study are listed in Supplementary Table S2.

### Folding of oligonucleotides

Guanine-rich oligonucleotides were denatured in 50 mM KCl for 3 min at 90°C, immediately transferred to 75°C and slowly cooled down to 10°C.

For the RNA duplexes used in the binding assays, 5′-^32^P-labeled oligomers were hybridized in water with 10 pmol of complementary oligomers by heating and slowly cooling the mixtures from 90°C to 4°C. The reaction mixtures were PAGE-purified in 12% native polyacrylamide gels to obtain double-stranded fractions free of single-stranded species.

### In-gel G-quadruplex staining

200 pmol of non-labeled oligomers were folded in 20 μL of 50 mM KCl, mixed with 20 μl of loading buffer, divided into two equal portions (20 μl) and separately loaded on a 12% polyacrylamide gel supplemented with 5% glycerol. The samples were arranged in duplicate so that after electrophoresis, the gel could be vertically cut into halves containing the same samples. One half was then stained in a solution of SYBR Gold (Thermo Fisher Scientific)/1×TBE for 20 min and scanned with Fujifilm FLA-5100 Fluorescent Image Analyzer at 473 nm. The other half was stained with *N*-methyl mesoporphyrin IX (NMM; Frontier 6 Scientific), as described previously (46). Briefly, NMM stock (5 mg/ml in 0.2 M HCl) was diluted in 20 mM Tris pH 7.6, 100 mM KCl, 1 mM EDTA to a final NMM concentration of 1 μg/ml (NMM staining solution). Next, the gel was incubated in the NMM staining solution for 20 min. Subsequently, the gel was scanned at 532 nm.

### *o*-BMVC labeling of oligonucleotides

In-house synthesized C6-amino-modified oligo dissolved in sodium phosphate buffer (0.1 M, pH 7.0) was combined with 5-fold molar excess of in-house made *o*-BMVC-C3-NHS ester in DMSO. The mixture was incubated at 37°C in the dark overnight. The oligo-*o*-BMVC-C3 conjugate was separated from salt and free *o*-BMVC-C3-NHS by 2% NaClO4/acetone precipitation. The purity and homogeneity of the oligo-*o*-BMVC-C3 conjugate were verified by denaturing gel electrophoresis.

### Preparation of the 5′–3′-end-ligated TER22

The 5′-^32^P-labeled TER22 was denatured in 100 mM KCl for 3 min at 90°C immediately transferred to 75°C and slowly cooled down to 10°C to adopt its native structure. The ligation reaction was carried out using T4 RNA Ligase 1 (Thermo Fisher Scientific) overnight at 17°C. RNA was purified using NucAway Spin Columns (Thermo Fisher Scientific) according to the manufacturers protocol and resuspended in water to the final concentration of approximately 10,000 cpm/µL (50 nM). To verify the efficiency of TER22 circularization, RNA (10,000 cpm) was treated with 0.5 U of alkaline phosphatase (Thermo Fisher Scientific) according to the manufacturers protocol and analysed by denaturing PAGE.

### RNA/DNA-protein binding assay

RNA/DNA-protein complex formation was analyzed using an electrophoretic mobility shift assay (EMSA). The reactions were carried out in 10-μL or 20-μL volumes. PPC or PPC variants (total protein concentration ranging from 0.1 to 400 nM (unless stated otherwise in the figure legend) were mixed with a trace amount of the 5′-^32^P-labeled RNA/DNA oligonucleotide (approximately 10,000 cpm) or *o*-BMVC-labeled oligonucleotide (from 7.5 to 30 µM, as indicated) and incubated in binding buffer (20 mM Tris-HCl, pH 7.4, 100 mM KCl) for 30 min (unless stated otherwise) on ice. The samples were separated in 5% or 10% non-denaturing polyacrylamide gels (as indicated) at 4°C for approximately 10 h at 7 V/cm in 1×TBE. For radiolabeled oligomers, gels were exposed to a phosphorimager plate, which was subsequently scanned with Fujifilm FLA-5100 Fluorescent Image Analyzer to visualize the bands. For *o*-BMVC-labeled oligomers, gels were first scanned with Fujifilm FLA-5100 Fluorescent Image Analyzer at 532 nm, then stained in a solution of SYBR Gold (Thermo Fisher Scientific)/1×TBE for 20 min and scanned again at 473 nm.

### hDicer cleavage inhibition assay

Dicer cleavage inhibition assay was performed in 10 µL reactions containing 20 mM Tris-HCl, pH 7.5, 100 mM KCl, and 2.5 mM MgCl2 buffer, 10 nM hDicer, 5′-^32^P-labeled pre-miRNA (10,000 cpm, approximately 5 nM) and the indicated oligonucleotide (0.1, 0.5, 2 µM). In addition, two control reactions were carried out: (i) a negative control (*C-*) with no enzyme and no inhibitor, to test the integrity of the substrate during the incubation time, and (ii) a positive control (*C+*) with enzyme but no inhibitor added. All samples were incubated at 37°C for 30 min. The reactions were halted by adding 1 volume of 8 M urea loading buffer and heating for 5 min at 95°C, and then separated in a 15% polyacrylamide gel with 7 M urea and 1×TBE.

In the time-course assay performed under the low-turnover conditions, a 10 µL reaction contained reaction buffer as described above, 10 nM hDicer, 5′-^32^P-labeled pre-miRNA (10,000 cpm, approximately 5 nM), and the indicated oligonucleotide (0.5 µM). Samples were incubated at 37°C for 30 min, 1.5 h, 3 h, 6 h, or 12 h. In addition, two negative control reactions were carried out to monitor the integrity of the substrate during the incubation time: (i) a sample frozen immediately after the mixture was prepared (*C0*) and (ii) a sample incubated at 37°C for 12 h (*C12*). After the incubation time, all of the samples were processed as described above.

For the high-turnover conditions, the time-course assay was performed similarly, except that a 10 µL reaction contained 0.5 nM hDicer, 25 nM 5′-^32^P-labeled pre-miRNA, and 50 nM oligonucleotide (as indicated).

### Gel imaging and data analysis

For binding and cleavage inhibition assays, the data were collected using Fujifilm FLA-5100 Fluorescent Image Analyzer and quantified using MultiGauge 3.0 software (Fujifilm). Data from binding experiments were fit to a saturation isotherm using Prism 8.2.1 (GraphPad). Dissociation constants were calculated using the equation for one site, specific binding: Y = Bmax*X/(Kd + X) and Bmax = 100% set as a constraint; Kd values together with the associated standard errors are reported in the figures. Diagrams presenting data from cleavage assays were prepared in Excel 2016. In the case of all diagrams, error bars represent SD values calculated based on three independent experiments.

### Production and purification of hDicer PPC cassette and the 3’-pocket and the 5’-pocket PPC variants

The PPC cDNA, which corresponds to the 316-amino acid (aa) sequence located between 753 and 1068 aa of hDicer, was amplified by PCR using a purchased plasmid encoding a complete *Homo sapiens* Dicer1 ribonuclease type III sequence (PubMed, NM_030621) (GeneCopoeia). In the case of PPC variants, we used plasmids encoding hDicer 3’-pocket double mutant (Y926F, R927A), and the 5’-pocket sextuple mutant (R778A, R780A, R811A, H982A, R986A, R993A) (23) (Addgene). PCR fragments were subsequently cloned into the pMCSG7 vector (courtesy of Laboratory of Protein Engineering, Institute of Bioorganic Chemistry, Polish Academy of Sciences), which allows introduction of a 6xHis tag at the N-terminus of the protein. hDicer PPC and the PPC variants were expressed in *E. coli* strain BL21Star (Thermo Fisher Scientific), in standard Luria-Bertani (LB) medium. Gene expression was induced with 0.4 mM IPTG and bacteria were cultured for 18 h at 18°C with shaking. The cell pellets were lysed and purified with Ni2+-Sepharose High-Performance beads (GE Healthcare) with imidazole gradient (0.02 M-1 M) in 25 mM HEPES (pH 8.0) (in the case of PPC), or 25 mM Tris (pH 8.0) (in the case of PPC variants), supplemented with 300 mM NaCl, 0.1% Triton X-100, and 5% glycerol. The protein purity was assessed by SDS-PAGE. Selected fractions containing homogeneous protein were concentrated using Amicon filters (Merck) in respective buffers (as described above) enriched with 40% glycerol, and stored at -20°C.

### Docking and molecular modeling

Docking of RNA and the protein was performed following a meta-approach using different docking methods to generate the docking poses, followed by the rescoring and selection of best poses (47). The hDicer (PDB entry 4NGF) and G-quadruplex (PDB entry 2M18) were docked using the following methods: 3dRPC/RPDOCK (48), ClusPro (49), HADDOCK (50), HDOCK (51), Hex (52), PatchDock (53) and ZDOCK (54). While 3dRPC and HADDOCK have modules to dock protein and RNA explicitly, ClusPro allows only the use of RNA as a receptor molecule. For the methods HDOCK, Hex, PatchDock, and ZDOCK the docking was performed both with protein as receptor and RNA as ligand and vice-versa. Additionally, the protein-RNA complex structures were generated with an in-house method SimRNP developed by the Bujnicki group, an extension of SimRNA (55) that enables flexible modeling of RNA structures as well as protein-RNA complexes (Michał Boniecki and. J.M.B., unpublished). Additional information on modeling are available in the Supplementary Methods. The simulation trajectory is presented in the Supplementary Fig. S7), and the representative frame is provided as PDB file (Supplementary File S1). PDB files for the reported models have been deposited in Figshare and can be accessed under the following URL: https://figshare.com/s/1dfdb3d2945d404e8065.

See Supplementary Information for additional methods.

## Acknowledgments

We would like to thank prof. Ryszard Kierzek for providing G-rich oligomers, dr. Anna Urbanowicz for a technical support in RNA-protein binding studies, and Life Science Editors for editing services. This work was supported by the National Science Centre, Poland [2016/22/E/NZ1/00422 to A.K-K., 2017/26/A/NZ1/01083 to J.M.B., 2017/01/X/ST5/00577 to D.B. and 2017/01/X/ST5/00589 to D.G.], the Polish Ministry of Science and Higher Education [KNOW program for years 2014-2018] and the IIMCB statutory funds [to J.M.B and C.N]. Computational analyses were performed using the resources of IIMCB, the Poznan Supercomputing and Networking Center at the Institute of Bioorganic Chemistry, Polish Academy of Sciences [grant 312], the Polish Grid Infrastructure [grant rnpmc] and the Interdisciplinary Centre for Mathematical and Computational Modelling at the University of Warsaw [grants G73-4 and GB76-30]. Funding for open access charge: [2016/22/E/NZ1/00422 to A.K-K.].

## Author Contributions

N.K., A.S., K.C., M.W., M.P., M.C.M., D.G., and D.B. performed experiments; C.N. and J.M.B. performed computational analyses and modeling; all authors analyzed the data, and interpreted the results; J.M.B., Z.G., M.F., and A.K.K. supervised and provided advice; N.K. and A.K.K. wrote the manuscript with input from all other authors.

