## Supplementary Information for "RNA and DNA G-quadruplexes bind to human Dicer and inhibit its activity"

**This PDF file includes:**

Supplementary Methods
Figures S1 to S7
Table S1
Table S2
Legend for Supplementary File S1
Legend for Dataset S1
Supplementary References

**Other supplementary materials for this manuscript include the following:**

Supplementary File S1
Dataset S1

### Supplementary Methods

#### $^{32}$P labeling of oligonucleotides

The 5′-end labeling was performed using 10 pmol of RNA/DNA, 1 μL γ-$^{32}$P-ATP (3000 Ci/mmol, Hartman Analytic GmbH) and 10 U T4 polynucleotide kinase (Thermo Fisher Scientific) in a final volume of 10 μL. After 10 min of the incubation at 37°C, the reaction was halted by adding 1 μL 0.5 M EDTA, pH 8.0. The radiolabeled oligonucleotides were PAGE-purified in 8% denaturing polyacrylamide gels and resuspended in water to a final concentration of approximately 10,000 cpm/μL.

#### RNA-RNA binding assay

RNA-RNA complex formation was analyzed using EMSA. The specific 5′-$^{32}$P-labeled pre-miRNA (10,000 cpm, approximately 5 nM) was mixed with the respective guanine-rich oligomer (100 pmol), and incubated in binding buffer (20 mM Tris-HCl, pH 7.5, 100 mM KCl) in a final volume of 10 μL for 30 min at 37°C. The samples were separated in 12% non-denaturing polyacrylamide gels supplemented with 5% glycerol at room temperature for approximately 6 h at 7 V/cm in 1×TBE.

#### hDicer production

Human Dicer was prepared as described previously (1). In brief, protein with a His-tag at the C terminus was expressed in Baculovirus Expression System (Life Technologies) and purified by $Ni^{2+}$ affinity chromatography (Ni-NTA Agarose, Qiagen) followed by ion-exchange chromatography (HiTrap Q HP, GE Healthcare). Non-reducing conditions were applied for both protein purification and storage.

#### Docking and molecular modeling

Docking of RNA and the protein was performed following a meta-approach using different docking methods to generate the docking poses, followed by the rescoring and selection of best poses (2). The hDicer (PDB entry 4NGF) and G-quadruplex (PDB entry 2M18) were docked using the following methods: 3dRPC/RPDOCK (3), ClusPro (4), HADDOCK (5), HDOCK (6), Hex (7), PatchDock (8) and ZDOCK (9). Additionally, the protein-RNA complex structures were generated with an in-house method SimRNP developed by the Bujnicki group (Michał Boniecki and. J.M.B., unpublished) as an extension of SimRNA (10).

From each of the methods, 100 top-scored docking poses (according to the internal scoring system of each method) were selected, resulting in a total of 1200 decoys. These selected top poses were re-scored using four different scoring functions: QUASI-RNP (11), ProSPR (S. Mukherjee, personal communication), 3dRPC/RPRANK (12), and ITScorePR (13). From each of these scoring trials, 100 top-scored poses were collected. These poses were superposed onto the coordinates of the cryo-EM structure of hDicer in complex with a pre-miRNA substrate (PDB ID: 5ZAL) (14) and inspected for clash with the pre-miRNA. The poses which clash with the pre-miRNA binding were selected.

Molecular dynamics simulations were performed with the selected poses using the Amber 18 package (15). The missing residues in the hDicer protein structure were built using the MODELLER (16) module in UCSF Chimera (17). The input structure for the simulation was prepared using tleap in a truncated octahedral box of 10 Å allowance using the TIP3P water model (18). The simulations were performed using the combination of the Amber ff14sb force field (19) for proteins and the χOL3 force field (20) for RNA. The structure was energy-minimized for 10,000 cycles with restraints, followed by 10,000 cycles without restraints. The minimized structures were subjected to heating, density equilibration and short runs of equilibration. The heating was done from 100 K to 300 K for 500 ps with restraints on the entire structure and the density equilibration was performed for 500 ps, also with restraints on the entire structure. The equilibration of the structures was run for four short rounds. The first three rounds of equilibration were run for 200 ps each with the main chain atoms constrained. The final round of equilibration was performed for 2 ns. The production run was run for 100 ns. The minimization was performed using Sander, and the subsequent steps were performed using the CUDA version of PMEMD available in the Amber package (21-23). The simulation trajectory (Supplementary Figure S7) was clustered and the representative frame is provided as PDB file in the Supplementary Data (Supplementary File S1).

Docking of G4T4G4 DNA (PDB ID: 1JPQ) and the hDicer protein (PDB ID: 4NGF) was performed using the following docking methods: ClusPro (4), HADDOCK (5), HDOCK (6), Hex (7), PatchDock (8) and ZDOCK (9). While HADDOCK have modules to dock protein and DNA explicitly, ClusPro allows only use of DNA as a receptor molecule. For the methods HDOCK, Hex, PatchDock and ZDOCK the docking was performed both with protein as receptor and DNA as ligand, and vice-versa. From each of these docking trials, 100 top scored docking poses (according to the internal scoring system of each method) were selected resulting in a total of 1000 decoys. These poses were superposed on to the cryo-EM structure of hDicer in complex with a pre-miRNA substrate (PDB ID: 5ZAL) (14) and inspected for clash with the pre-miRNA substrate. The poses that clash with the pre-miRNA binding were selected.

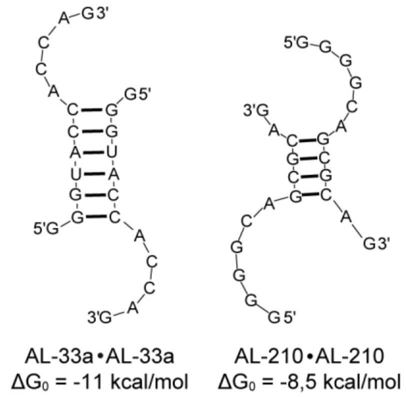


**Fig. S1 AL-33a and AL-210 display potential for homodimerization.** Schematic representation of the secondary structure predicted for AL-33 (*left*) and AL-210 (*right*).

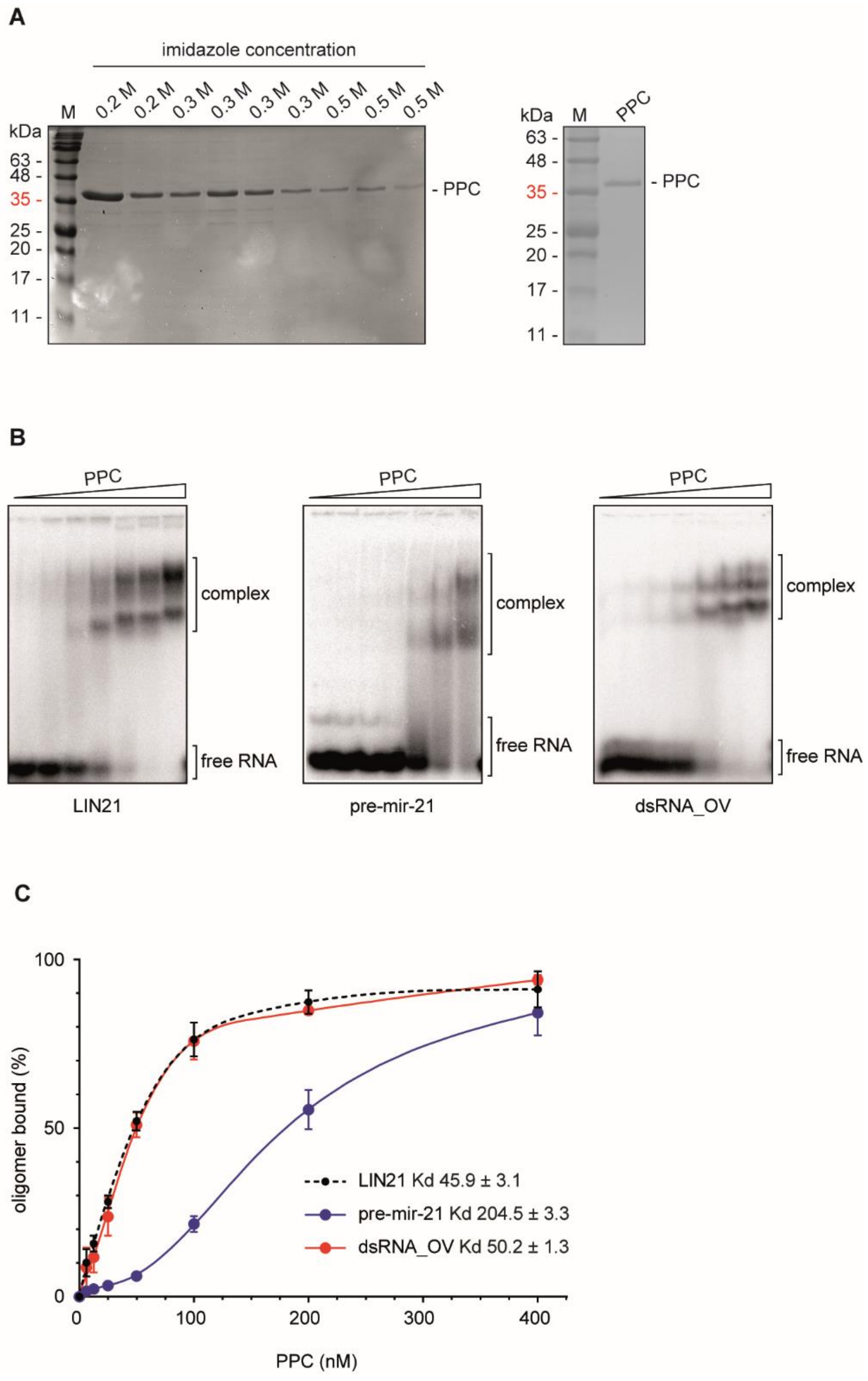


**Fig. S2 Biochemical characteristics of hDicer PPC cassette. (A)** SDS-PAGE analysis of the fractions collected after purification of PPC using $Ni^{2+}$- affinity chromatography (*left*) and the final PPC preparation used in the further assays (*right*). The analyzed protein fractions were washed out with buffers containing different imidazole concentration as indicated. Proteins were detected by Coomassie brilliant blue staining. M – protein size marker (11-245 kDa, EURx). (**B**) EMSAs with PPC and 5′-$^{32}$P-labeled ssRNA: 21-nt LIN21, 58-nt pre-mir-21 or dsRNA: 19-bp dsRNA_OV with 2-nt 3′-overhangs. Triangles represent increasing amount of PPC (6.25, 12.5, 25, 50, 100, 200, 400 nM). Free RNA and the RNA in complex with PPC are indicated. (**C**) The Kd values of RNA-PPC complexes were determined from the binding isotherms by curve fitting using nonlinear regression. Error bars represent SD from three separate experiments.

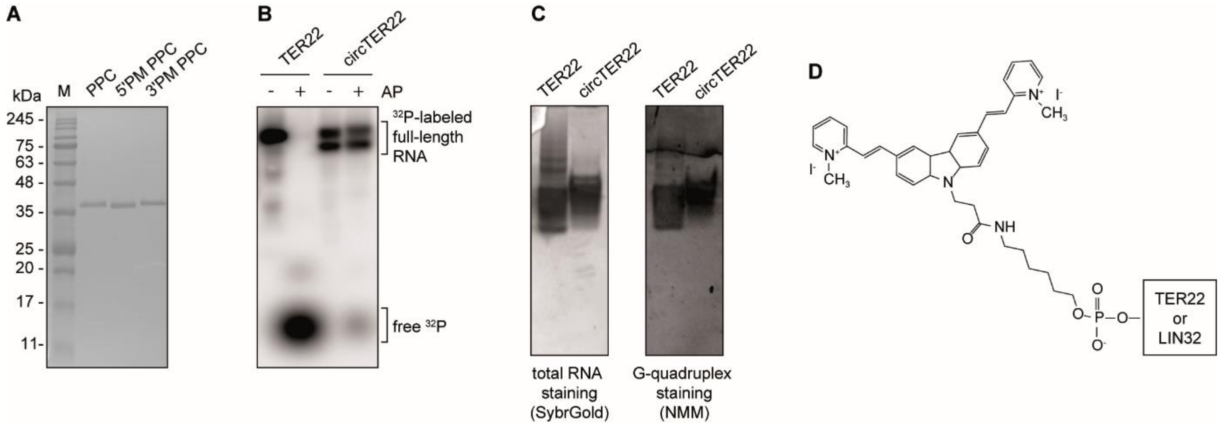


**Fig. S3 Characterization of hDicer PPC cassette preparations and G-quadruplexes used for binding assays. (A)** SDS-PAGE analysis of wild-type PPC and PPC 3′- and 5′-pocket mutants (PM). M – protein size marker (11-245 kDa, EURx). **(B)** Verification of the efficiency of TER22 circularization. Shrimp alkaline phosphatase (AP) dephosphorylates 5′-$^{32}$P-labeled TER22 with free ends but not after successful 5′-3′ end ligation. **(C)** Native PAGE analysis of linear or circularized TER22. Gels were treated with nucleic acid stain SYBR Gold (*left*) or G-quadruplex-specific dye, NMM (*right*). **(D)** Schematic representation of the structure of *o*-BMVC-C3-labeled oligomers.

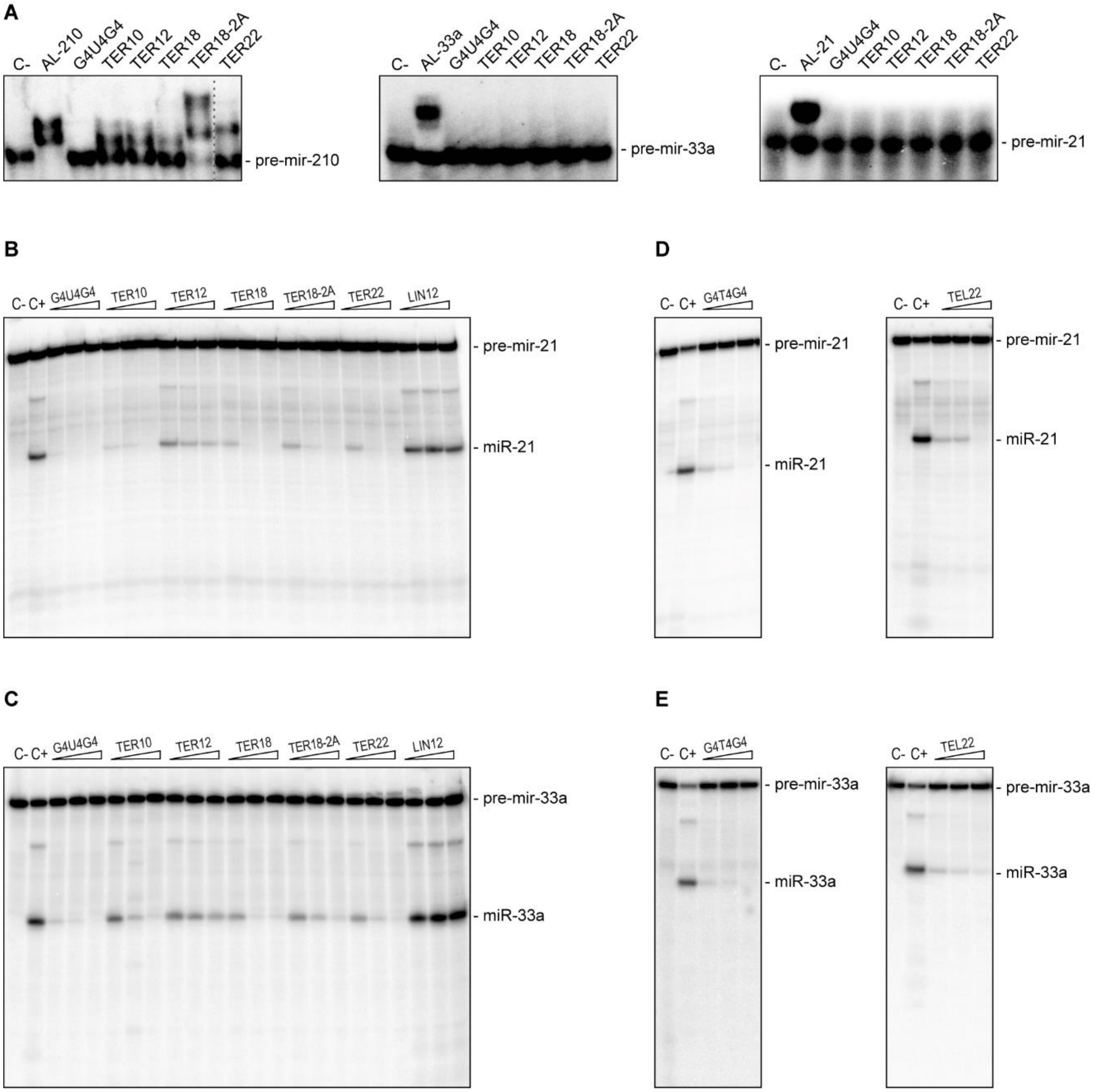


**Fig. S4 Characterization of the effect exerted by G-quadruplexes on pre-miRNA cleavage by Dicer.** **(A)** Native PAGE analysis of 5′-$^{32}$P-labeled (10,000 cpm, approximately 5 nM) pre-mir-210, pre-mir-33a, pre-mir-21 incubated with a a complementary 12-nt long RNA (AL-210, AL-33 or Al-21, respectively) and a set of RNA G-quadruplexes (10 μM). The position of a band corresponding to free pre-miRNA is indicated. Pre-mir-33a and pre-mir-21, as pre-miRNAs non-interacting with tested G-rich oligomers, were used as substrates in Dicer cleavage inhibition assays. *C-* a sample with pre-miRNA only. **(B, C)** Representative results of PAGE analysis used to quantify the effect of RNA G-quadruplexes (G4U4G4, TER10, TER12, TER18, TER18-2A, TER22) on the cleavage of pre-mir-21 (B) or pre-mir-33a (C) by hDicer. *C-* a sample with no protein, nor inhibitor added, *C+* a sample with hDicer, without inhibitor. Triangles represent an increasing amount (0.1, 0.5, 2 μM) of the indicated oligomer. LIN12 – a control 12-mer not adopting a G-quadruplex structure. **(D, E)** Representative results of PAGE analysis used to quantify the effect of DNA G-quadruplexes (G4T4G4, TEL22) on the cleavage of pre-mir-21 (D) or pre-mir-33a (E) by hDicer. *C-* a sample with no protein, nor inhibitor added, *C+* a sample with hDicer, without inhibitor. Reactions were carried out for 30 min at 37°C. Triangles represent increasing amount of the indicated oligomer (0.1, 0.5, 2 μM).

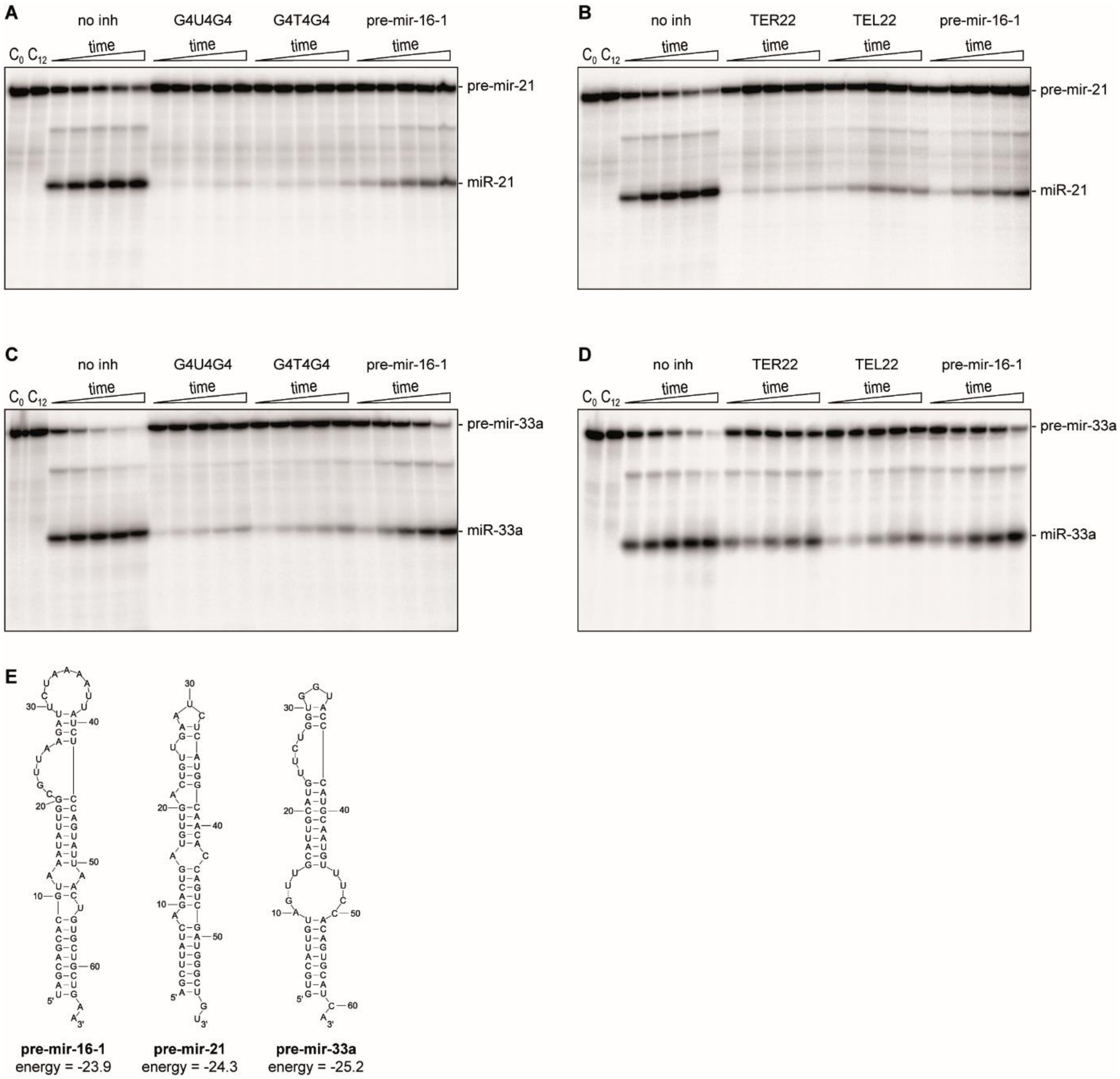


**Fig. S5 The time course for inhibition of pre-miRNA cleavage by Dicer mediated by G-quadruplexes under low-turnover conditions. (A-D)** Representative PAGE results used for quantitative analysis of the time course of hDicer (10 nM) inhibition by RNA and DNA G-quadruplexes (500 nM) in reactions with 5′-$^{32}$P-labeled pre-miRNA (10,000 cpm, approximately 5 nM): pre-mir-21 (A, B) or pre-mir-33a (C, D). Reactions were carried out at 37°C for: 30 min, 1.5 h, 3 h, 6 h, 12 h (increasing incubation time is indicated by a triangle). $C_0$, $C_{12}$ controls with no oligomer, nor hDicer added, stopped immediately after reaction assembly or after 12 h of incubation, respectively. **(E)** The lowest free energy structures predicted by RNAstructure Fold algorithm for pre-miRNAs used in the assay. The free energy values are expressed in kcal/mol.

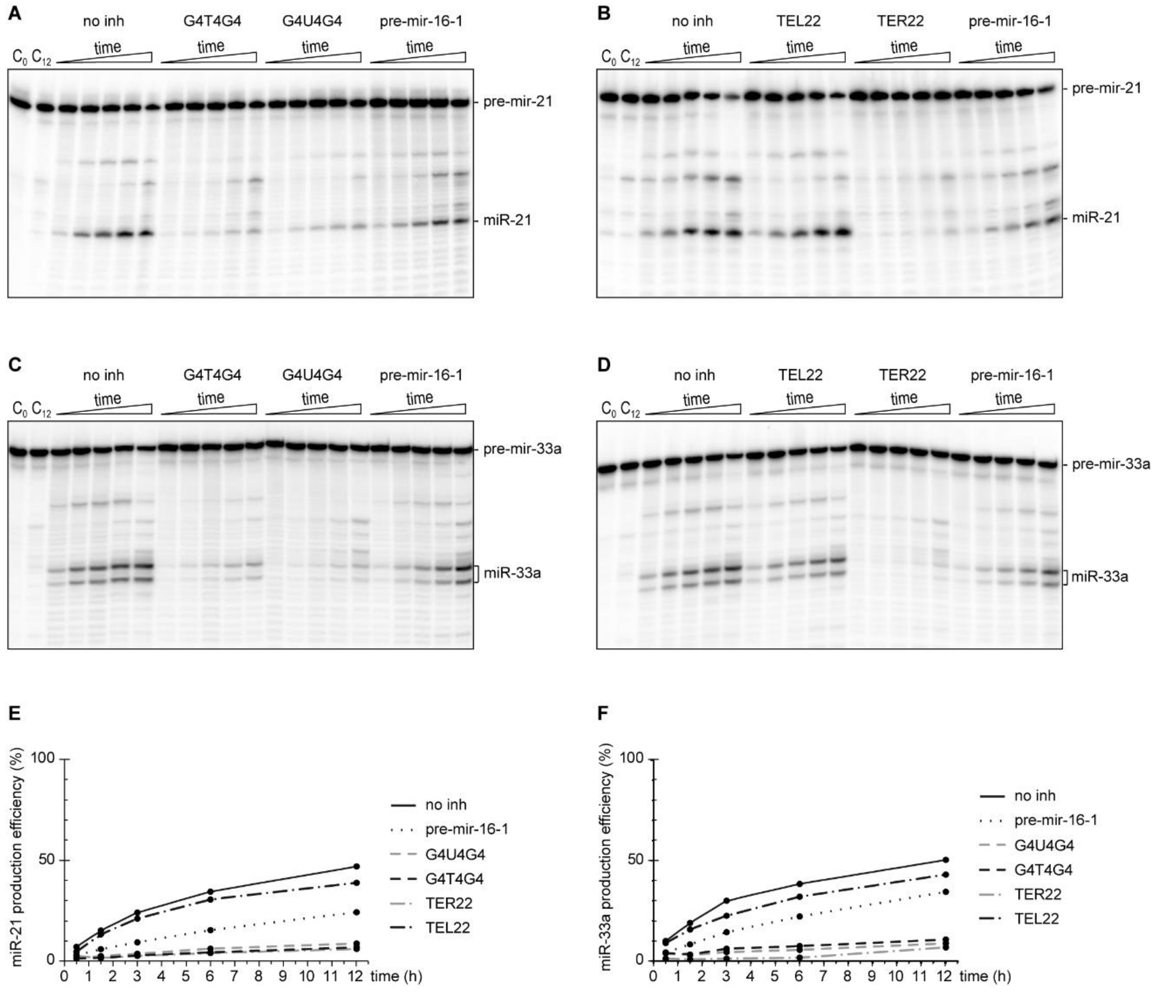


**Fig. S6 The time course for inhibition of pre-miRNA cleavage by Dicer mediated by G-quadruplexes under high-turnover conditions. (A-D)** PAGE results used for quantitative analysis of the time course of hDicer (0.5 nM) inhibition by RNA and DNA G-quadruplexes (50 nM) in reactions with 5′-$^{32}$P-labeled pre-miRNA (100,000 cpm, approximately 25 nM): pre-mir-21 (A, B) or pre-mir-33a (C, D). Reactions were carried out at 37°C for: 30 min, 1.5 h, 3 h, 6 h, 12 h (increasing incubation time is indicated by a triangle). $C_0$, $C_{12}$ controls with no oligomer, nor hDicer added, stopped immediately after reaction assembly or after 12 h of incubation, respectively. **(E-F)** Quantitative analysis of the time course of hDicer inhibition by G-quadruplexes in reactions with pre-mir-21 (E) or pre-mir-33a (F).

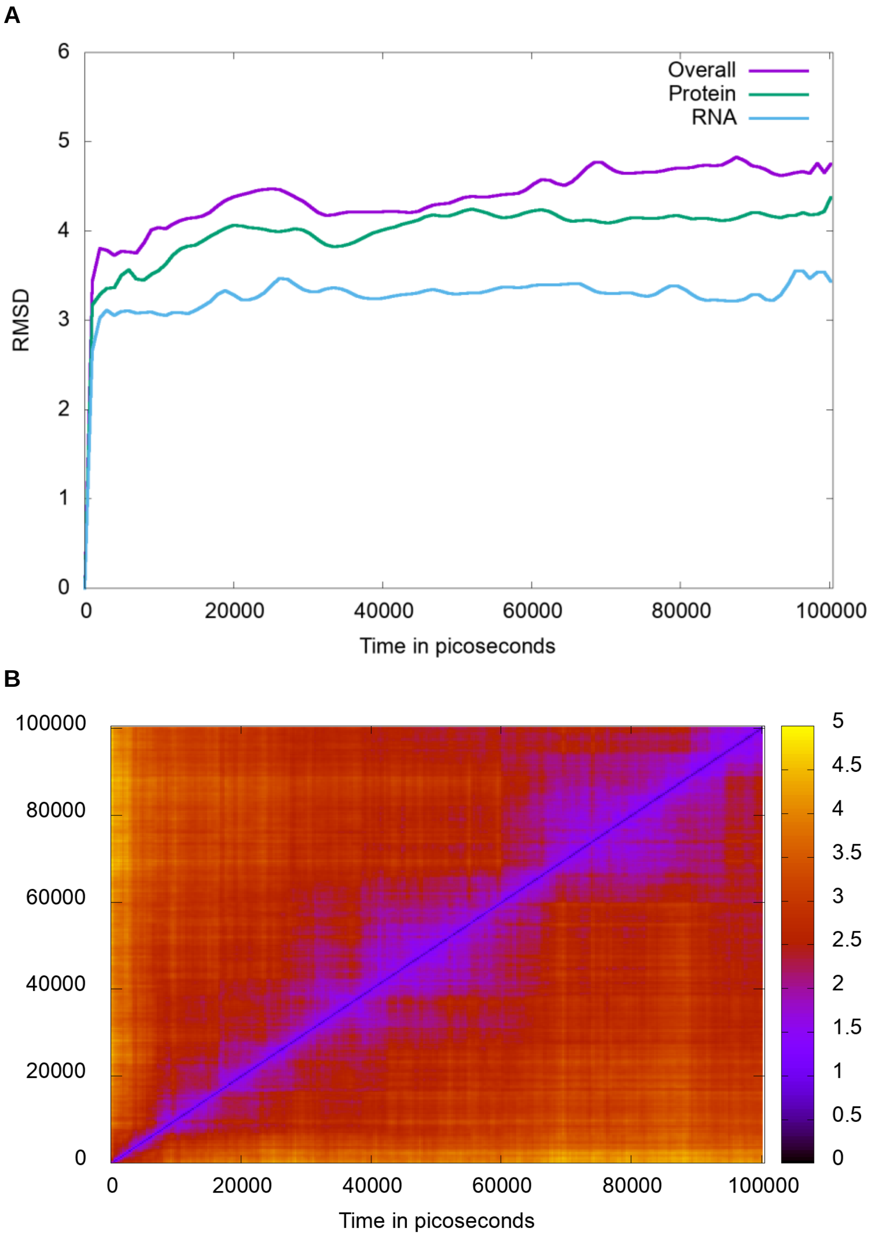


**Fig. S7 Molecular dynamics simulations of hDicer and G-quadruplex complex**. **(A)** The root mean square deviation (RMSD) of the frames with respect to the starting frame of the simulations remains stable throughout the simulations. **(B)** The heat map of pairwise RMSD of different frames during the MD simulations; points are colored based on the RMSD.

**Table S1. Characteristics of RNA and DNA quadruplexes used in the study.**

| **Name** | **Sequence** | **PDB ID** | **Type** | **Topology** | **Reference** |
|---|---|---|---|---|---|
| TER10 | r(GGGUUAGGGU) | 2M18 | dimer of bimolecular quadruplexes | parallel | (24, 25) |
| TER12 | r[(UAGGGU)$_2$] | 2KBP | bimolecular | parallel | (25) |
| TER18 | r(GGGUUGCGGAGGGUGGGC) | - | monomolecular or dimer of monomolecular quadruplexes | parallel | (26) |
| TER18-2A | r(AAGGGUUGCGGAGGGUGGGC) | - | monomolecular | parallel | (26) |
| TER22 | r[AGGG(UUAGGG)$_3$] | - | monomolecular | parallel | (27) |
| G4U4G4 | r(GGGGUUUUGGGG) | - | bi- or tetramolecular | parallel | (28) |
| QU14 | r(GGAGGUUUUGGAGG) | 1MY9 | dimer of monomolecular quadruplexes | parallel | (29) |
| TEL22 | d[AGGG(TTAGGG)$_3$] | - | monomolecular | mixture of parallel/antiparallel | (30) |
| G4T4G4 | d(GGGGTTTTGGGG) | 1JPQ | bimolecular | antiparallel | (31, 32) |

**Table S2. Sequences of all oligonucleotides used in the binding and cleavage assays.**

| **Name** | **Sequence (5′ → 3′)** |
|---|---|
| AL-16-1 | GAAUCUUAACGC |
| AL-21 | AUGAGAUUCAAC |
| AL-33a | GGGUACCACCAG |
| AL-210 | GGGGCAGCGCAG |
| LIN21* | UCGAAGUAUUCCGCGUACGUG |
| LIN32 | GUGCAUUGUAGUUGCAUUGCAUGUUCUGGUCA |
| dsRNA_OV | CGUACGCGGAAUACUUCGAAA |
| TER10 | GGGUUAGGGU |
| TER12 | UAGGGUUAGGGU |
| TER18 | GGGUUGCGGAGGGUGGGC |
| TER18-2A | AAGGGUUGCGGAGGGUGGGC |
| TER22 | AGGGUUAGGGUUAGGGUUAGGG |
| TEL22 | AGGGTTAGGGTTAGGGTTAGGG |
| G4U4G4 | GGGGUUUUGGGG |
| G4T4G4 | GGGGTTTTGGGG |
| QU14 | GGAGGUUUUGGAGG |
| pre-mir-16-1 | UAGCAGCACGUAAAUAUUGGCGUUAAGAUUCUAAAAUUAUCUCCAGUAUUAACUGUGC UGCUGAA |
| pre-mir-21 | AGCUUAUCAGACUGAUGUUGACUGUUGAAUCUCAUGGCAACACCAGUCGAUGGGCUGU |
| pre-mir-33a | GUGCAUUGUAGUUGCAUUGCAUGUUCUGGUGGUACCCAUGCAAUGUUUCCACAGUGCAU CA |
| pre-mir-210 | GCCCCUGCCCACCGCACACUGCGCUGCCCCAGACCCACUGUGCGUGUGACAGCGGCUG |

** LIN21 also serves as a complementary strand to form dsRNA with dsRNA_OV*

**Supplementary File S1 (separate file). A representative frame from the molecular dynamics simulation of hDicer and G-quadruplex complex.**

**Dataset S1 (separate file). Guanine-rich sequences identified in the transcriptome-wide map of hDicer targets.**
